# EBEx: an Ensemble-Based Explainable Framework for Gene Calling in Heterogeneous Diseases

**DOI:** 10.64898/2026.03.12.710464

**Authors:** Iria Pose-Lagoa, Beatriz Urda-García, Nuria Olvera, Jon Sánchez-Valle, Rosa Faner, Alfonso Valencia, José Carbonell-Caballero

## Abstract

Complex and clinically heterogeneous diseases pose significant challenges for gene prioritisation and patient stratification, as relevant genes often show weak or context-specific signals and transcriptomic datasets are limited in size. These limitations hinder the discovery of robust molecular signatures using traditional case-control approaches and motivate computational pipelines capable of capturing molecular diversity. Here, we present an explainable ensemble-based AI pipeline to prioritise disease-relevant genes from transcriptomic data, using Chronic Obstructive Pulmonary Disease (COPD) as a use case. To retain biologically relevant interactors obscured by molecular heterogeneity, the framework integrates data-driven signals with curated COPD-related gene sets, further expanded through network-based prioritisation and supported by molecular interactions. Gene relevance is evaluated via aggregated explainability scores across multiple classifier configurations to ensure robust candidate selection. The final set comprised < 8% of evaluated genes, ∼ 62% arising from network-based expansion, substantially reducing dimensionality while preserving biological heterogeneity. Beyond case-control classification, the approach identified candidate genes and molecular subgroups associated with specific clinical features, capturing patient-level heterogeneity. The prioritised genes recapitulated key disease-related processes, including immune responses and extracellular matrix degradation, and highlighted additional associations like the enrichment of the IL-4 and IL-13 signalling pathway, which is of clinical interest given ongoing biologic developments targeting these axes. Our pipeline outperformed existing methods in discriminating COPD from controls, and the final gene list was validated in independent cohorts. Implemented as a scalable and reusable R package, this framework facilitates the study of molecular heterogeneity in complex diseases like COPD, supporting advances in diagnosis and precision medicine.

**Availability and implementation:** EBEx code and tutorials can be found in: https://iposelag.github.io/EBEx/

## Introduction

Understanding the molecular basis of complex diseases is challenging due to the interaction of genetic, environmental, and lifestyle risk factors across the lifespan, leading to diverse clinical and molecular profiles. Many complex conditions, such as Chronic Obstructive Pulmonary Disease (COPD), comprise poorly defined or overlapping patient subgroups that standard clinical classifications fail to capture. COPD is a prevalent chronic respiratory disorder characterized by persistent airflow limitation, with smoking as the primary risk factor (1). Although post-bronchodilator FEV1 thresholds define clinical disease severity, individuals with comparable airflow limitation can present different molecular alterations and clinical trajectories (2; 3). This heterogeneity complicates case-control classification, particularly in mild disease stages where clinical definitions may not reflect molecular variability.

Transcriptomic technologies offer opportunities to apply Artificial Intelligence (AI) techniques and investigate such heterogeneity (4; 5). However, transcriptomic studies are typically constrained by limited cohort sizes relative to the number of measured genes. This high-dimensional, low-sample-size problem reduce signal to noise ratio, destabilises feature selection, and limits the applicability of complex models such as Deep Learning, which are prone to overfitting under these conditions (6; 7). In COPD, previous microarray-based studies have combined Differential Expression Analysis (DEA) with regression models, network-based module detection, or single-classifier frameworks to derive disease-associated signatures (8; 9; 10). While informative, these approaches often rely on univariate filtering, prioritise a single predictive model, or evaluate feature importance within a specific modelling framework, potentially limiting robustness in heterogeneous disease contexts.

A key challenge of univariate strategies for dimensionality reduction is their inability to capture multivariate gene interactions (11). Genes with weak individual effects may nonetheless contribute to disease mechanisms through coordinated activity within biological networks. Integrating multivariate feature selection strategies, such as minimum Redundancy Maximum Relevance (mRMR), with systems biology approaches enables identification of genes within disease-relevant molecular contexts. Complementarily, model-agnostic explainability techniques such as SHAP (SHappley Additive exPlanations) quantify gene contributions across different model architectures at both global and individual levels, facilitating cross-model comparison and improving interpretability (12).

We therefore hypothesise that no single classifier is universally optimal for modelling a heterogeneous disease such as COPD in a case-control framework. Instead, we propose an ensemble-based pipeline to identify a diverse and robust set of COPD-associated genes beyond those detected by classical approaches such as DEA. To test this hypothesis, we 1) formulated COPD molecular analysis as a supervised prediction task (cases vs. controls) and trained well-established classifiers on gene lists derived from transcriptomics data analysis and prior biological knowledge. 2) We expanded these lists using a systems-biology perspective that combines network-based prioritisation and molecular interaction databases to retrieve new potential candidate genes that operate in the same biological pathways. 3) We then applied a model-agnostic explainability method to quantify gene contributions across scenarios. 4) Finally, we derive a procedure that works with aggregated SHAP values to obtain a unified gene relevance score, enabling the identification of candidate genes that consistently contribute to disease prediction. Based on ranked scores and classification, a final procedure associates genes with clinically relevant phenotypes, opening new possibilities for the interpretation of the molecular basis of diseases and specific conditions (e.g. emphysema).

## Methods

### Feature Selection Approaches

To identify genes informative for COPD prediction, we applied multiple feature selection strategies to the training data and evaluated the resulting sets in subsequent classification tasks (Fig. 1, Fig. 2b).

**Figure 1.**
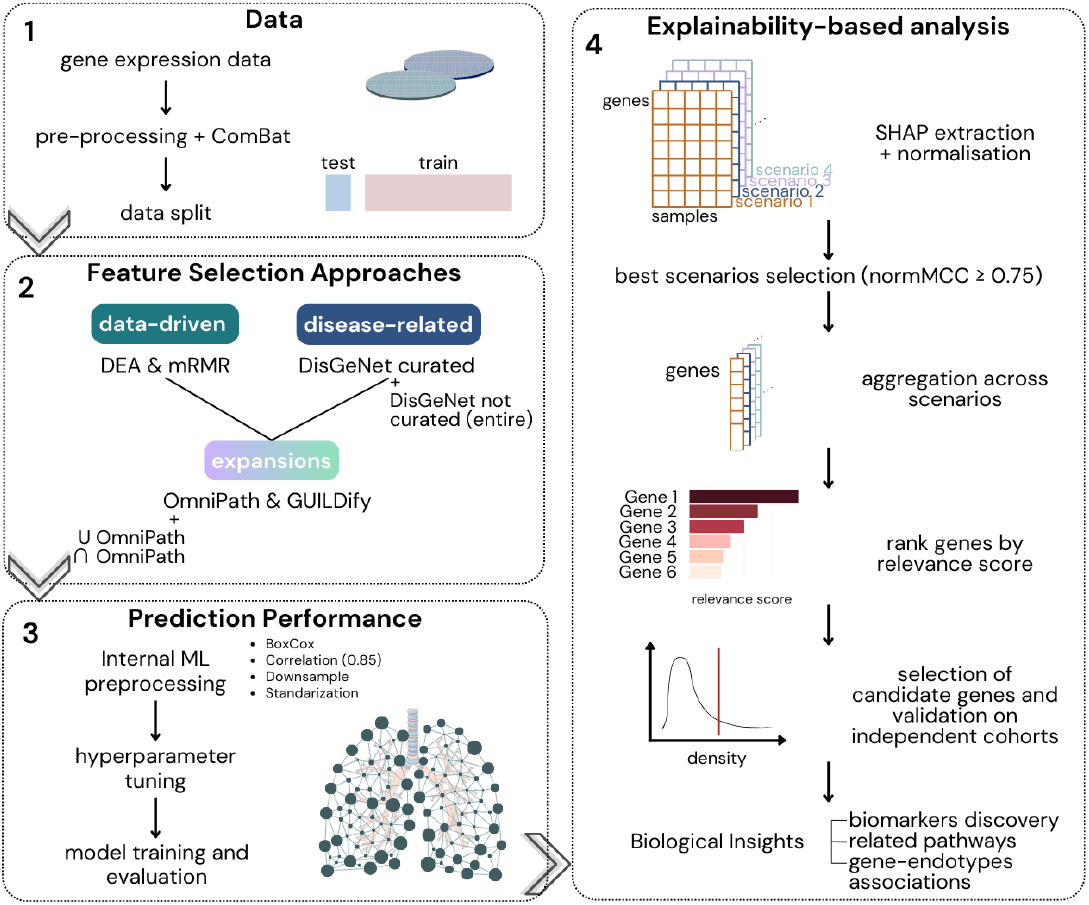
Flowchart of the pipeline. (1) Two transcriptomic platforms were preprocessed and integrated using ComBat (Note S1), followed by partitioning into training and independent test sets. (2) Three complementary feature-selection strategies were evaluated: (i) data-driven genes derived from intrinsic expression patterns; (ii) curated COPD-related genes retrieved from DisGeNET; and (iii) network-based expansions of these seed lists using OmniPath and GUILDify. (3) A standardised preprocessing recipe was applied within the training set. Model hyperparameters were optimised via Bayesian optimisation, and final classifiers were trained on the full training data, assessed by 10-fold cross-validation, and evaluated on the independent test set. (4) Model explainability analyses were performed across all scenarios to compute aggregated gene relevance scores. These scores were also aggregated across scenarios and ranked to define the final candidate gene set, which was subsequently validated in independent cohorts. The final stage translates predictive signals into biological insight, identifying biomarkers, regulatory mechanisms, and molecular COPD endotypes.

**Figure 2.**
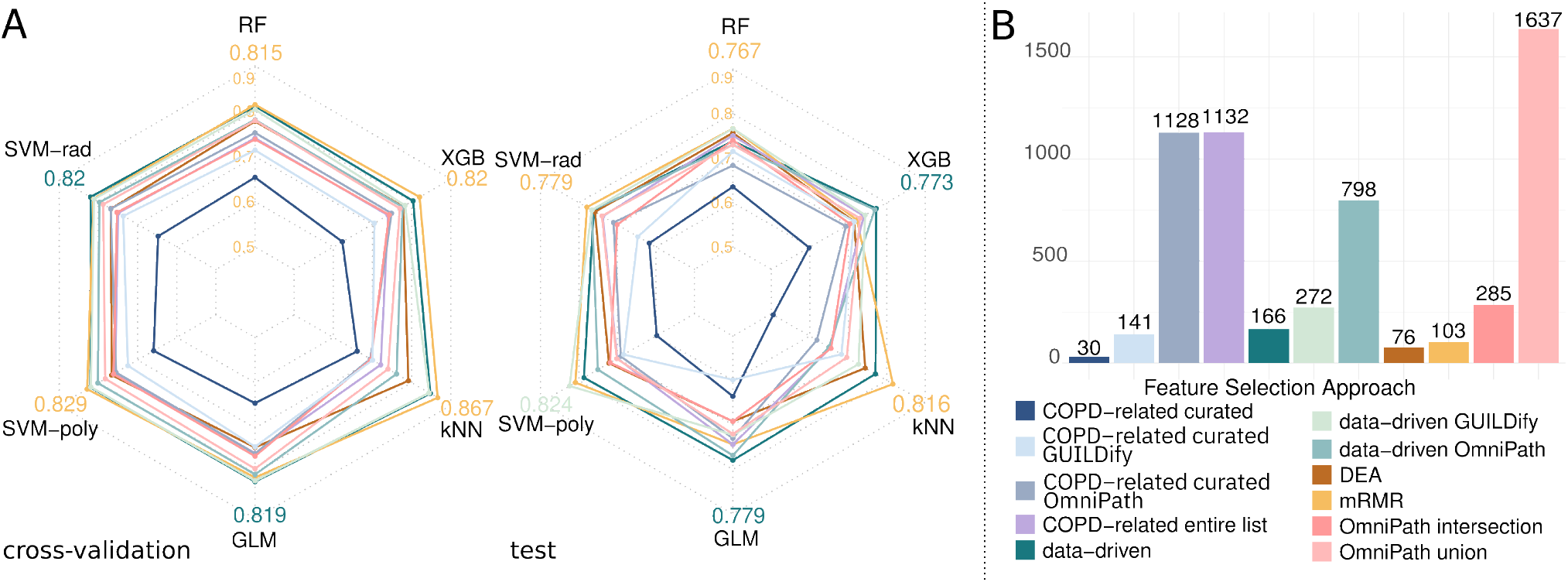
Prediction performance across all scenarios. (A) normMCC scores for each ML model (vertices), with colours indicating the input gene set. The left panel shows cross-validation results; the right panel shows test results. (B) Barplot displaying the number of genes in each input list.

#### Data-driven identification of COPD-discriminative genes

Genes driven by intrinsic data characteristics were identified through two complementary approaches. First, DEA between cases and controls was performed using the limma framework (11), adjusting for platform and sex covariates. Genes were considered significantly differentially expressed (DEA genes) if they met False Discovery Rate (FDR) ≤ 0.01 and Fold Change (FC) ≥ 1.5. Second, we applied mRMR, a multivariate feature selection method that prioritises genes maximising class discrimination while minimising redundancy (13). Genes were ranked according to their contribution to case–control separation, and optimal gene sets were determined based on classification performance and statistical significance against null distributions (Note S2).

#### Disease-related gene identification of COPD

To incorporate prior biological knowledge, we retrieved disease-related genes from DisGeNET using the term *“Chronic Obstructive Airway Disease (CUI: C0024117)”* (14). Two gene sets were defined: expert-curated associations (COPD-related curated) and the full set of reported associations (COPD-related entire).

#### Biological network-based expansions of seed gene lists

Data-driven and COPD-related curated gene sets were expanded using two complementary network-based approaches. First, physical interaction partners were identified using OmniPath (15), retrieving both directed and undirected interactions with the seed genes. The union and intersection of the resulting expanded lists were computed and interactions with a curation effort score below 1 were excluded to ensure data quality. Second, gene prioritisation was performed using GUILDify (16), which integrates heterogeneous biological data and network-based prioritisation algorithms. Following the authors’ recommendations (17), we selected the top 1% highest-scoring non-seed genes identified by the NetScore, NetZcore, and NetShort methods.

#### Literature validation

To assess the association between seed genes and COPD, we queried PubMed to identify prior publications in which each gene was mentioned in association with COPD (Note S3).

### Classifiers Training and Hyperparameter Optimisation

COPD analysis was formulated as a binary classification task distinguishing COPD patients from controls, with COPD as the positive class. The dataset was split into training (75%) and test (25%) sets. Given class imbalance, model optimisation was guided by normalised Matthew’s Correlation Coefficient (normMCC). Prior to training, model-specific preprocessing steps were implemented according to the requirements of each classifier. To capture complementary learning strategies, we evaluated multiple classifiers: Random Forest (RF) (18), SVM with radial and polynomial kernels (SVM-rad and SVM-poly, respectively) (19), Generalised Linear Models (GLM) (20), k-Nearest Neighbours (kNN) (21), and XGBoost (XGB) (22). Hyperparameters were optimised using grid search and Bayesian optimisation, with model selection based on mean normMCC under 10-fold cross-validation (Fig. 1, Note S4).

To assess whether observed performance exceeded random expectations, we generated 1000 null models by randomly shuffling gene identities while preserving input set size and compared model accuracy against the null distribution (Fig. S8).

### Explainability-based relevance score for gene prioritisation

Model-agnostic explainability was applied to quantify gene contributions to classification performance across modelling scenarios. SHAP values were computed for each sample and gene (23). To enable comparison across classifiers and input lists, SHAP values were normalised per sample (normSHAP) within each scenario and aggregated by gene, capturing their overall predictive importance independently of effect direction. To ensure robustness, only classifier–gene list combinations achieving high predictive performance (normMCC *>* 0.75 in both cross-validation and independent test sets) were considered. normSHAP values from these selected scenarios were aggregated to derive a unified relevance score for each gene, reflecting its maximal contribution across modelling configurations. A threshold was subsequently defined by modelling the distribution of relevance scores and identifying the point at which the density and its local derivatives stabilised (yielding a threshold of 0.0109), thereby delineating the upper tail of the importance distribution (Fig. S9). An alternative clustering-based strategy was used as validation (Note S5).

Functional enrichment of the candidate genes was performed with enrichR (24), leveraging pathway and ontology annotations from Reactome (25), GO (26), and KEGG (27) (FDR ≤ 0.05).

### Patient heterogeneity and gene characterisation

Aggregated explainability scores were available at the patient–gene level, enabling the evaluation of associations between candidate genes and clinical variables across COPD samples using appropriate statistical tests (Wilcoxon rank-sum (28), Student’s t-test (29), or Pearson’s chi-squared test (30); adjusted p-value ≤ 0.05) (Note S6). Patient-level heterogeneity was characterised through hierarchical clustering based on aggregated normSHAP values or gene expression profiles (Euclidean distance). Clinical enrichment analyses were then repeated within each subgroup to identify subgroup-specific molecular and clinical patterns (Note S7). Cross-tissue expression patterns and lung cell-type specificity of candidate genes were assessed using publicly available GTEx bulk RNA-seq data and Human Cell Atlas (Note S8).

### Validation analysis based on independent cohorts

Model robustness was evaluated using independent lung tissue cohorts generated by microarray and RNA-seq technologies (see Data availability section). For ML validation, batch effects between the initial dataset and each external cohort were corrected before evaluation. Models were retrained on the initial dataset using only candidate genes shared with each external cohort and tested on the corresponding external dataset after quantile normalisation. Additionally, we assessed whether candidate genes were enriched among DEA genes (evaluating multiple FDR and log2FC thresholds) identified independently in each cohort. To determine whether observed overlaps exceeded random expectations, a Monte Carlo permutation-based analysis was performed by comparing observed overlaps to a null distribution generated from randomly sampled gene sets of identical DEA genes size.

## Results

In this study, we analysed the transcriptome of 301 lung tissue samples (Note S1). COPD patients had a mean age of 64.96 ± 9.68 years and were slightly predominantly male (57.92%), while the controls had a mean age of 64.25 ± 11.05 years with balanced sex distribution (Table 1).

**Table 1.** Summary statistics of phenotypic variables. Absolute frequencies and mean ± sd for categorical and continuous data, respectively.

|  | COPD | Controls |
| --- | --- | --- |
| N of samples | 202 | 99 |
| Age | 64.96 ± 9.67 | 64.25 ± 11.05 |
| Sex | 117 Males (57.92%)<br>85 Females (42.08%) | 47 Males<br>47 Females |
| Smoker | 14 Current<br>177 Ever<br>8 Never<br>3 NA | 2 Current<br>60 Ever<br>29 Never<br>8 NA |
| Predicted FEV1 (% ref)* | 54.67 ± 21.99 | 99.91 ± 12.71 |
| Predicted FVC* | 80.49 ± 18.42 | 97.33 ± 13.17 |
| Predicted DLCO | 56.03 ± 22.87 | 83.75 ± 16.77 |
| Emphysema (f-950) | 15.37 ± 16.86 | 0.61 ± 1.00 |

DLCO: Diffusing Capacity of the Lung for Carbon monoxide; FEV: Forced Expiratory Volume in 1 second; FVC: Forced Vital Capacity; \*post Bronchodilator test; NA: Missing values

### Complementary strategies reveal genes missed by classical approaches

To identify genes discriminating COPD from controls, we applied complementary feature-selection strategies designed to capture signals that may be missed by classical approaches such as DEA or curated disease-related genes. We evaluated three sources of information: (i) data-driven genes derived from intrinsic expression patterns, (ii) curated COPD-related genes from DisGeNET (14), and (iii) network-based expansions using OmniPath (15) and GUILDify (16) (Fig. 1).

The data-driven strategy combined classical DEA results and the features ranked by the mRMR algorithm (see Methods). DEA identified 76 genes (50 overexpressed, 26 underexpressed). However, since biologically relevant signals may not reach classical DEA thresholds, we complemented DEA with mRMR feature selection (31). Model performance declined beyond 100 top-ranked mRMR genes, consistent with null-distribution analyses identifying 103 meaningful features (Note S2). Combining DEA and mRMR yielded 166 genes with minimal overlap (13 genes), highlighting methodological complementarity (Fig. S5). Literature review showed that 27.1% of data-driven genes had prior COPD associations, while others (e.g., *ROR1, TGFB2*) had limited direct evidence but plausible mechanistic links, suggesting recovery of both established and underexplored markers (Note S3).

To incorporate prior knowledge, we retrieved COPD-related genes from the DisGeNET database (14), defining both curated COPD-related set (30 genes) and an entire COPD-related set (1,132 genes). Curated COPD-related set showed strong literature support but limited overlap with data-driven genes (Note S3, Fig. S5). Only *MMP1* (3.33%) overlapped with the curated core. This discrepancy may reflect tissue specificity, as curated databases integrate associations across multiple tissues and contexts. These results indicate that highly cited disease genes are not necessarily optimal discriminators in lung transcriptomic data, highlighting the complementarity between curated knowledge and data-driven discovery.

Finally, to account for pathway-level effects beyond individual genes, both data-driven and curated seed sets were expanded using biological network information (Fig. 2b; Methods). These network-based approaches broadened the molecular landscape, recovering other candidates that are less studied but may still be relevant to the disease.

### Comparative performance across classifier-gene list combinations

We evaluated Bayesian-optimised ML classifiers to predict COPD cases and controls using the generated gene lists. Figure 2a (Table S2) shows normMCC performance across all classifier-gene list combinations, with consistent trends for accuracy (Fig. S11).

Gene lists derived from data-driven strategies, particularly mRMR and data-driven sets, consistently achieved the highest normMCC values in cross-validation and testing. The curated COPD core showed the lowest performance, whereas expanding it to the disease-related entire list substantially improved normMCC, highlighting the limitations of small curated sets. Network-based expansions generally outperformed their corresponding seed lists in cross-validation, though testing showed more variability, suggesting that expansions help capture complementary disease-relevant signals. Notably, comparison against null models confirmed that only data-driven list achieved accuracies significantly above random expectation, whereas COPD-related curated set did not (adjusted p-value ≤ 0.01; Fig. S8, Methods).

Rather than identifying a single optimal scenario, these findings suggest that multiple scenarios capture complementary predictive signals, motivating their integration in a subsequent explainability-based analysis.

### Scenario-specific explainability reveals complementary COPD markers

To evaluate how different modelling scenarios prioritise predictive genes, we compared normSHAP-based feature importance rankings across all classifier–gene list combinations. This analysis aimed to determine whether the same genes consistently ranked highly or whether importance depended on the specific scenario.

Comparison of the top 30 normSHAP-ranked genes per scenario revealed limited overlap across classifiers, even within the same input list. This indicates that different modelling strategies capture complementary predictive signals, consistent with COPD heterogeneity. Among gene lists derived from intrinsic transcriptomic properties, only six genes appeared in more than two-thirds of the 30 scenarios. In contrast, COPD-related lists showed lower overlap, with only *D8A* and *TGFBR3* recovered in half of the scenarios. Notably, network-based expansions of COPD-related seed genes introduced additional high-importance genes, indicating that literature-derived seeds capture only part of the predictive landscape. Conversely, data-driven lists showed greater stability across scenarios, with lower added contribution from their expansions (Fig. S12).

These results highlight that different modelling scenarios capture complementary sets of predictive genes, supporting their combination to identify robust COPD-associated genes.

### Final candidate gene set identifies canonical and extended markers

To derive a final set of COPD candidate genes, we aggregated normSHAP values across the best-performing modelling scenarios, selecting genes in the tail of this score relevance distribution (aggregated normSHAP ≥ 0.0109, Methods). This yielded 172 genes, representing *<* 8% of all evaluated genes (Fig. S9, Table S3). A comparable gene set was obtained using an alternative clustering-based approach, confirming the robustness of the selection (Note S5).

Candidates were classified depending on whether they could be recovered by classical strategies: *canonical* (identified by DEA or the curated COPD-related list) or *extended* (identified exclusively by mRMR, network expansions, or the disease-related entire list). Overall, 38.4% were *canonical* and 61.6% *extended*, indicating that most highly predictive genes would not be recovered by classical univariate or curated strategies alone.

The candidate genes originate from diverse inputs, highlighting the complementarity of the initial feature-selection strategies (Fig. 3b, Fig. S13). DEA contributes 36 (20.93%) *canonical* genes, while the curated COPD-related list contributes one gene, with one additional gene recovered by both sources. Among the *extended* genes, 53 (30.81%) were selected from mRMR, 31 (18.02%) from all intersecting expansions, 8 (4.65%) from data-driven expansions, and 9 (5.23%) from disease-related expansions. Together, these results show that data-driven strategies and their network-based approaches account for the majority of candidate genes.

**Figure 3.**
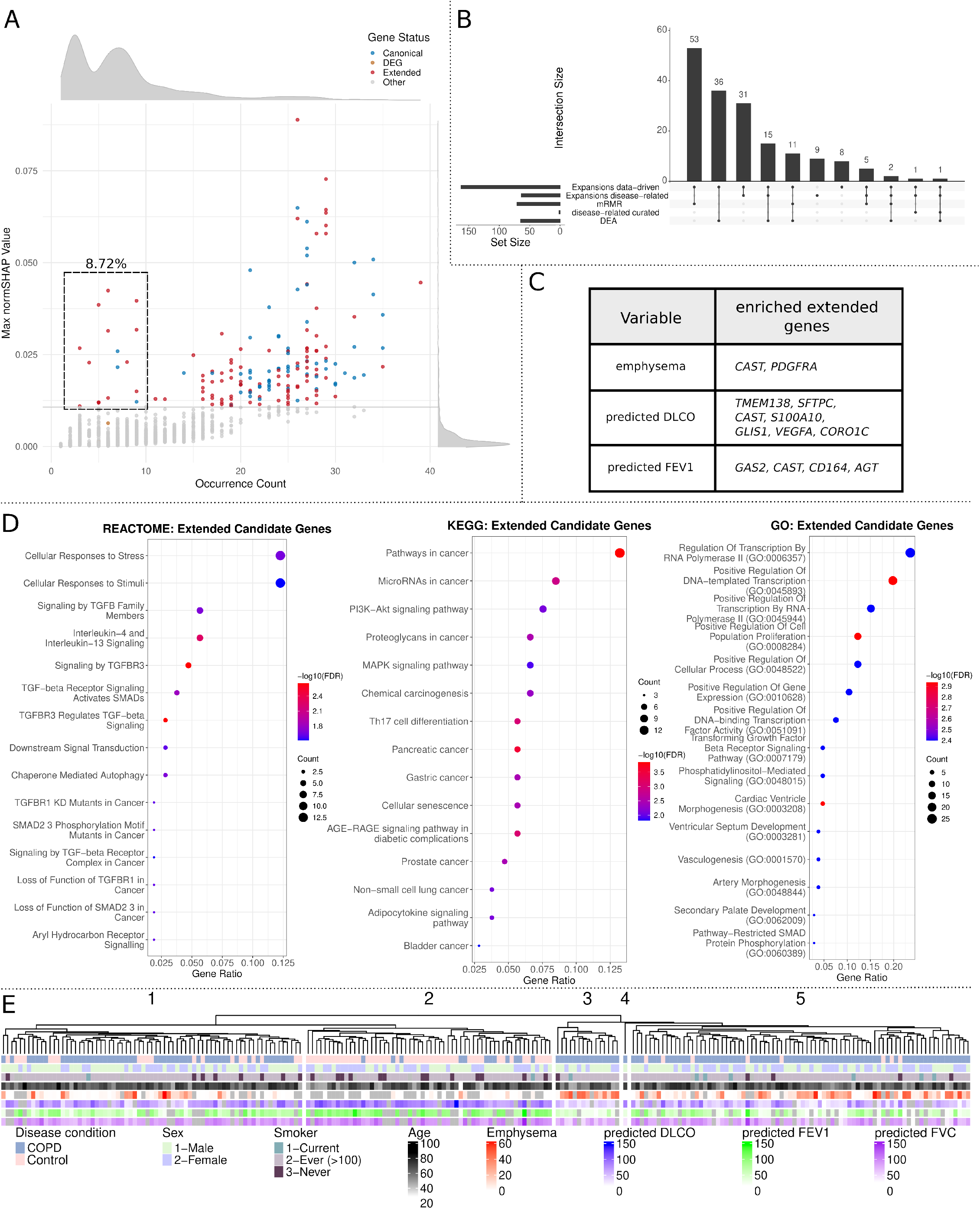
Identification and characterisation of candidate genes. (A) Scatter plot showing the relationship between the maximum aggregated normSHAP value and the number of scenarios in which each gene appears. Candidate genes are those above horizontal line (y = 0.0109). Genes are coloured by classification: *canonical* (blue), *extended* (red), and genes identified exclusively by DEA but not retained in the final candidate set. (B) UpSet plot summarising the origin of candidate genes, grouped by their source gene list. (C) Table listing enriched (*extended*) genes across clinical variables. (D) Top 20 enriched pathways of *extended* genes based on Reactome, KEGG and GO analysis. (E) Hierarchical clustering of samples based on candidate gene expression, with top annotation representing clinical variables.

Figure S14 shows that 62.21% of the candidate genes fall outside conventional DEA FC thresholds, and 23.26% of our candidate genes have neither significant FDR values nor large FC, indicating that they carry predictive signals not captured by classical univariate statistics. This behaviour suggests that these genes participate in multivariate patterns or interact with other features in ways that may remain undetected by the applied DEA and curated resources.

Moreover, we observed a positive correlation between the relevance score value and the number of scenarios in which it appeared (*R* = 0.69, *p* ≤ 2.2 × 10^−16^; Fig. 3a, Fig. S15), indicating that consistently selected genes tend to have higher predictive importance. However, 8.72% high-importance genes appeared only in specific combinations, revealing context-dependent markers detectable under certain classifiers-gene list interactions.

### Biological interpretation and patient molecular heterogeneity

Our final candidate set included both well-known COPD-associated genes and novel potential biomarkers. Several candidates have also been implicated in related respiratory diseases and comorbid conditions, supporting shared inflammatory and remodelling mechanisms across chronic lung disorders (Note S9).

Functional enrichment analyses showed that *canonical* genes were mainly associated with immune dysregulation and extracellular matrix organisation, consistent with established COPD pathogenesis (Methods, Fig. S16) (32). In contrast, *extended* genes were enriched in pathways linked to structural remodelling and regulatory signalling, including TGF-*β* signalling (33), SMAD phosphorylation (34), and epithelial-to-mesenchymal transition (35). Additionally, several KEGG pathways emerged, such as PI3K–AKT, and those related to hypoxia, focal adhesion or cancer. Notably, enrichment also highlighted pathways corresponding to recently proposed therapeutic targets, including Aryl Hydrocarbon Receptor signalling and Interleukin-4 and Interleukin-13 signalling (Fig. 3d) (36; 37).

Clinical association analyses of the aggregated scores revealed genes (*extended*) associated with lung function measures, including lung function impairment and emphysema severity (Fig. 3c). Associated genes include markers of alveolar epithelial integrity (*SFTPC*), regulators of vascular remodelling and angiogenesis (*VEGFA*), and mediators of fibroblast activation (*PDGFRA*). Additional genes point to immune and inflammatory regulation (e.g., *S100A10*) and cilia-related epithelial maintenance (*TMEM138*), supporting the contribution of structural, vascular, and immune components to lung function decline (Note S6).

Cross-tissue expression analysis showed that *canonical* genes were predominantly lung-enriched, whereas *extended* genes displayed broader tissue distributions, suggesting potential systemic involvement. A subset of candidates showed expression in both lung and blood, supporting their potential as minimally invasive biomarkers. Moreover, certain candidates exhibit enriched expression in lung and other organs (breast, uterus, liver or brain), which may reflect sex-specific effects, comorbidities, or environmental exposures such as smoking (Note S8) (38; 39; 40; 41).

Unsupervised clustering based on the expression of the candidate genes stratified patients into COPD and controls, and identified distinct molecular subgroups. These included a severe airflow-limitation endotype (cluster 3), a heterogeneous cluster (cluster 5), and a healthier and non-smoking subgroup (cluster 2), reflecting the intrinsic molecular complexity of the disease (Fig. 3e, Note S7).

### External validation in lung cohorts and comparison with other studies

Candidate genes were validated across multiple independent lung tissue cohorts. In a combined microarray dataset integrating GSE8581 and GSE37768 (61 samples), 154 candidate genes were retained among 13, 933 shared genes, and ML models showed consistent above-random performance under strict external testing. Similar results were observed in the GSE69818–GSE103174 cohort (153 COPD and 37 control), where 157 candidate genes were present among 13, 855 shared genes (Table S6). In GSE76925 cohort enriched for advanced COPD stages (102 cases, 35 controls), candidate genes were retained, and unsupervised clustering clearly separated COPD and controls (Fig. S23). DEA (FDR ≤ 0.05, log2FC ≥ 0.5) identified 53 genes, with 11 candidate genes overlapping significantly beyond random expectation (even using different threshold elections). Moreover, validation using RNA-seq data revealed a significant overlap between candidate genes and DEA results (FDR ≤ 0.05, log2FC ≥ 0.5), indicating robustness across distinct transcriptomic technologies (Table S7).

Comparison with previously published microarray-based ML models (8; 9; 10) showed that our best-performing scenarios achieved high sensitivity (up to 0.97) and high specificity (up to 0.86) in cross-validation, surpassing the penalised logistic regression approach reported by Gohari et al. On the internal test set, performances remain robust (normMCC up to 0.81), and results were stable when using the final candidate gene set, supporting the robustness of the selected genes (Table S8).

## Discussion

Human complex diseases such as COPD are characterised by multifactorial aetiology and pronounced patient heterogeneity, complicating the identification of underlying molecular mechanisms and robust biomarkers. Although clinical stratification relies on phenotypic measures such as lung function or symptom severity, these metrics do not fully capture the molecular diversity underlying disease progression (1; 2; 42). These challenges underscore the need for computational frameworks capable of uncovering biologically meaningful patterns and identifying actionable targets.

To address these challenges, we developed an ML-based pipeline to identify candidate genes that capture both global transcriptional alterations distinguishing COPD from controls and context-specific signals in patient subgroups. Our approach integrates multiple well-established classifiers, such as RF and SVM, with systematically optimised hyperparameters to improve performance and reproducibility. The modular design of the pipeline also allows future integration of additional classifiers as new algorithms emerge, ensuring adaptability. To further enhance sensitivity, classifiers were trained on complementary gene sets, encompassing both curated disease-related genes and data-driven candidates that exhibit differential behaviour between controls and cases. Crucially, we expanded these lists using network-based gene prioritisation information from GUILDify and curated molecular interactions from OmniPath, enabling the recovery of genes that may play important roles in disease but could be overlooked by traditional univariate approaches such as DEA, which are often limited by COPD heterogeneity. Candidate selection was restricted to scenarios achieving high normMCC values, ensuring robust predictive models. To ensure specificity, explainability scores were aggregated across all experiments to quantify the overall contribution of each of the 2, 426 evaluated genes, yielding a concise (*<* 8%) yet informative portrait of their contribution across patients. An analytical threshold based on the distribution of these relevance scores allowed us to select a final set of 172 highly relevant candidate genes. Importantly, this list includes both *canonical* genes identified by classical approaches and *extended* genes uncovered through network expansions. The predominance of *extended* genes highlights the added value of combining complementary feature-selection strategies, ensemble-modelling, and model-agnostic explainability metrics.

*Canonical* candidate genes recapitulate known COPD mechanisms, including immune and inflammatory response and extracellular matrix degradation, validating our approach. In contrast, *extended* genes highlighted additional processes such as transcriptional regulation and tissue morphogenesis, suggesting broad regulatory shifts. Some of the identified genes, were related to injured/inflamed cell types such as alveolar cells and fibroblasts. Notably, enrichment of IL-3 and IL-4/IL-13 pathways aligns with current efforts to develop biologic therapies for these axes (36; 37). Moreover, the presence of cancer-related pathways and overlap with genes reported in lung cancer, asthma, and idiopathic pulmonary fibrosis further suggests convergent molecular responses to chronic lung injury (such as smoking) across chronic respiratory diseases and related comorbidities (43).

Cross-tissue analyses further reinforced biological relevance. While *canonical* genes tend to show lung-enriched expression consistent with established COPD biology, *extended* genes display more diverse expression profiles, including lung-specific, multi-tissue, blood-associated and even liver or brain expression patterns. Genes expressed in both lung and blood highlights their potential as non-invasive biomarkers, whereas genes shared with other organs may reflect systemic effects or comorbidities. The identification of genes with sex-biased tissue expression may help explain sex-specific differences observed in COPD susceptibility and progression (38; 44). Similarly, brain-expressed genes may reflect how persistent lung inflammation and oxidative stress in COPD can affect the brain, altering the blood-brain barrier and neuroendocrine regulation (45).

Importantly, we found that a binary case-control classification through the lens of different ML classifiers and candidate gene lists analysis allowed us to identify genes associated with specific clinical characteristics, thereby informing patient stratification. DLCO and FEV1 showed strong associations with *extended* candidate genes involved in alveolar lung tissue damage (e.g. *SFTPC, VEGFA*) and immune response (e.g. *S100A10*). Although no enrichment was observed for smoking status –possibly due to limited pack-year data– the candidate list includes genes such as *AHR* and *CYP1B1* that mediate responses to cigarette smoke toxicants. This may also suggests that our approach prioritises genes linked to functional and structural changes, while general exposure effects may be less distinguishable within this cohort.

Beyond individual gene–trait associations, clustering based on candidates’ gene expression separated COPD patients from controls and revealed distinct molecular endotypes associated with clinically relevant features, including airflow limitation, emphysema severity, and gas exchange impairment. Rather than focusing solely on disease status, this approach highlights molecular subgroups within COPD. At the same time, the presence of clusters without clear clinical enrichment reflects the intrinsic molecular heterogeneity of COPD and supports the coexistence of multiple disease mechanisms within clinically similar patients (3).

The robustness of the candidate gene set was supported by validation across multiple independent lung cohorts and transcriptomic platforms. Despite differences in study design, disease severity, and technology, candidate genes consistently showed significant overlap with DEA genes and supported disease-control separation in unsupervised analyses. Across the different modelling scenarios explored, the best-performing models for each evaluation metric outperformed previously published transcriptomic approaches. Notably, models trained on the final candidate gene set achieved improved or competitive performance—particularly in balanced metrics—compared with previously published transcriptomic models, supporting stability and generalisability (8; 9; 10).

However, several limitations should be acknowledged. First, our analysis relies on bulk lung tissue, potentially masking cell-type-specific effects, thus single-cell or spatial transcriptomics may contribute to refining the cellular and subpopulation-specific context of candidate genes. Second, clinical metadata were limited, restricting deeper phenotype–genotype associations. In addition, the associations identified here are predictive rather than causal, and integrating causal inference frameworks and longitudinal data represents an important direction for future research. For instance, emerging interventional or causal Shapley approaches, which incorporate underlying causal structure, could help distinguish potential disease candidates from correlated signals and further strengthen interpretability (46).

In conclusion, combining data-driven feature selection, network-based expansion, and explainable ML provides a robust framework to identify biologically meaningful candidate genes in COPD. By capturing both *canonical* and *extended* molecular signals, this approach uncovers established and underexplored mechanisms, refines patient stratification, and reveals gene-enodtype associations that were not previously apparent, highlighting potential biomarkers and therapeutic targets. Beyond COPD, this framework offers a generalizable strategy to extract mechanistic insight from heterogeneous transcriptomic data, bridging predictive modelling with biological interpretation. Implemented as an openly available R package, it provides a scalable and reproducible blueprint for studying other complex diseases characterised by molecular and clinical heterogeneity.

## Supporting information

Full suplementary text file including supplementary text, figures and tables

## Data availability

Microarray and RNA-seq datasets are publicly available in the Gene Expression Omnibus (https://www.ncbi.nlm.nih.gov/geo/) under accession numbers GSE47460, GSE76925, GSE8581, GSE37768, GSE69818, GSE103174, and GSE57148 (47; 48; 49; 50).

