## Supplementary material for "EBEx: an Ensemble-Based Explainable Framework for Gene Calling in Heterogeneous Diseases": Full suplementary text file including supplementary text, figures and tables

#### CONTENTS

1 Supplementary Notes 2
2 Supplementary Figures 5
3 Supplementary Tables 27

### 1. SUPPLEMENTARY NOTES

#### S1 - Data Preprocessing

Microarray gene expression samples from the Lung Tissue Research Consortium (LTR; GSE47460) were obtained from the Gene Expression Omnibus (https://www.ncbi.nlm.nih.gov/geo/) [1]. Samples were generated using two Agilent platforms (Agilent-014850 Whole Human Genome Microarray 4x44K G4112F and Agilent-028004 SurePrint G3 Human GE 8x60K), which were processed separately using a dedicated Agilent microarray pipeline (Fig. S1). Background correction and between-array quantile normalisation were first applied, followed by the removal of control probes and probes lacking GeneSymbol annotation. Low-quality samples (misannotated samples and outliers) and lowly expressed probes (expression below half the mean count of the minority group in the DEA) were subsequently filtered out (Fig. S1). The two platforms were then merged by retaining only common genes and correcting for platform-specific batch effects using the ComBat empirical Bayes framework [2]. Finally, 301 samples remained; 67% of the subjects were COPD patients, while the remaining 33% were marked as at risk (controls) (Table 1).

#### S2 - mRMR Feature Selection and threshold determination

Unlike univariate DEA, mRMR prioritises features that are both highly informative for class separation and minimally redundant with previously selected genes, using a Mutual Information Difference criterion on discretised expression data. The C/C++ implementation from Hachuan Peng's laboratory was used [3]. This method produced a ranked list of the top 500 features with an associated score. To define an optimal gene set size, we evaluated classification performance across multiple classifiers trained on the top $k$ genes from the mRMR ranking, with $k$ ranging from 10 to 500 (Fig. S2). Moreover, we conducted a bootstrapping analysis by shuffling sample disease labels and running 1000 arbitrary mRMR, thereby generating a null distribution of the feature score. Based on this distribution, we defined a score threshold of $a = 0.031$ ($P(x \in [a, 1]) \leq 0.01$) and retained genes above this threshold (Fig. S3).

#### S3- Literature Validation of seed-genes and COPD associations.

To assess the association between seed genes and COPD, we performed a literature search in PubMed using the easyPubMed R package [4]. The following query was used for searching publications in which each gene was mentioned in association with COPD:

```
"((GeneSymbol) AND ((Chronic Obstructive Lung Disease[MeSH Terms]) OR
(Chronic Obstructive Pulmonary Diseases[MeSH Terms]) OR (COAD[MeSH Terms]) OR
(COPD[MeSH Terms]) OR (Chronic Obstructive Airway Disease[MeSH Terms]) OR
(Chronic Obstructive Pulmonary Disease[MeSH Terms]) OR(Airflow Obstruction, Chronic[MeSH Terms]) OR
(Airflow Obstructions, Chronic[MeSH Terms]) OR (Chronic Airflow Obstructions[MeSH Terms]) OR
(Chronic Airflow Obstruction[MeSH Terms])))"
```

Fig. S4 shows that a subset of data-driven genes (27.1%) had been previously linked to COPD (e.g. *PLA2G7*, *FFG*, *PPARGC1A*), supporting the validity of our data-driven feature selection approach [5–7]. At the same time, mRMR also prioritised genes with limited prior evidence in COPD, such as *ROR1*, a regulator of Wnt signalling that in turn has been associated to lower alveolar epithelial progenitor capacity in COPD or *TGFB2*, whose nearby genetic variants have been associated with emphysema [8, 9]. These genes may provide links between established pathways that influence COPD through more indirect or underexplored mechanisms.

In contrast, curated COPD-related genes exhibited the strongest literature support, with an average of 107.6 publications per gene and some genes, such as *TNF* and *IL6*, exceeding 900 publications linked to the known systemic inflammatory profile of COPD patients.

Additionally, we see that the overlap with the data-driven list was limited (Fig. S4 and Fig. S5). Only *MMP1* (3.33%) from the curated gene set was identified in the intersection (increasing to 22 when considering the entire disease-related list), which is a well-stablished mediator of airway inflammation and epithelial dysfunction [10]. Thus, this discrepancy suggests that highly cited disease genes are not necessarily those that best discriminate cases from controls in a specific transcriptomic context, highlighting the complementary nature of curated knowledge and data-driven discovery.

#### S4-Classification details

All models were trained to perform a binary classification task distinguishing COPD patients from control subjects, with COPD defined as the positive class. The dataset was internally split into training (75%) and test (25%) sets, with the training set comprising 151 COPD samples and 74 controls. Given the resulting class imbalance, model optimisation was guided by Matthew's Correlation Coefficient (MCC), a robust metric for binary classification that yields high values only when predictions are accurate across all outcome categories [11]. For interpretability, we consider the normalised MCC metric, $\text{normMCC} = \frac{\text{MCC}+1}{2}$, as the performance metric to evaluate the methods alongside accuracy.

Prior to model training, the training data underwent a series of preprocessing steps. Features were rescaled using Box–Cox transformation and standardisation to approximate normality. Highly correlated features were filtered using a

Pearson correlation threshold of 0.85 to reduce multicollinearity. Class imbalance was addressed through down-sampling of the majority class. Model-specific preprocessing pipelines were implemented according to the requirements of each classifier (Fig. S6).

###### *Classifier implementation details*

Classifier models were implemented using the ranger, kernlab, glmnet, kNN, and xgboost engines, with additional support from tidyverse-based R packages [12].

###### *Tuning methodologies*

Hyperparameter optimisation was performed using two strategies: grid search and Bayesian optimisation (Table S1, Fig. S6). The grid search optimisation approach used a regular grid with a predefined set of parameter values. Alternatively, the Bayesian optimisation used a Gaussian Process model to iteratively propose improved hyperparameter configurations, using an expected improvement acquisition function to assign scores to these candidate solutions. Bayesian optimisation was initialised with six parameter combinations, allowed up to 25 iterations, and stopped if there were 15 iterations without improvement. As Bayesian optimisation outperformed grid search in both predictive performance and computational efficiency (Fig. S7), all reported models were tuned using the Bayesian strategy. Optimal hyperparameters were selected based on normMCC using a resampling technique. Specifically, we employed a 10-fold cross-validation strategy. For each configuration, one fold was used for evaluation while the remaining folds were used for training, and the configuration with the highest mean normMCC was retained (Fig. S6).

##### S5 - Explainability-based relevance score computation and candidate genes threshold determination

We used SHAP to assess the contribution of individual features to the predictions of the classifiers [13]. SHAP enables local model interpretation by quantifying how each feature influences the prediction for a specific observation. For each observation in our dataset, SHAP values were calculated by averaging feature contributions across different feature orderings. All calculations were conducted using the R package DALEXtra [14, 15]. To standardise comparisons across classifiers and input configurations, SHAP values were normalised per sample within each scenario using the formula:

$$\text{normSHAP}_{i,j} = \frac{\text{SHAP}_{i,j}}{\sum_{k=1}^{n} |SHAP_{ik}|}, i = 1, ..., m, j = 1, ..., n$$

where $i$ represents samples ($m$ = total number of samples) and $j$ represents features ($n$ = total number of features). normSHAP values were subsequently aggregated by gene to compute average absolute contributions, thereby capturing each gene's overall importance independently of the direction of its effect. Candidate genes for COPD classification were identified by retaining only combinations of input gene lists and classifiers achieving a normMCC greater than 0.75 in both cross-validation and test evaluations. normSHAP values derived from these selected combinations were aggregated by gene into a unified table, enabling genes to be ranked according to their maximum normSHAP value across all scenarios (relevance score).

To identify a threshold for selecting final candidate genes based on normSHAP values, we perform a density-based analytical approach (Fig. S9). The distribution of aggregated normSHAP values was first modelled using kernel density estimation to capture the overall shape of the data. A sliding window approach was then applied along the normSHAP value range to quantify changes in density values and their slopes. For each window along the normSHAP value range, differences in density values and slope were computed. These differences were analysed to pinpoint the region where the density curve and its derivatives stabilised, yielding a normSHAP value threshold of 0.0109. Genes with normSHAP values above this threshold were identified as candidates for further analysis (Table S3). To further validate this threshold, an independent clustering-based approach was applied (Fig. S10). Aggregated normSHAP values were clustered using one-dimensional Gaussian mixture models [16] to identify latent subpopulations of genes with similar importance scores. The number of clusters was selected using the Bayesian Information Criterion (BIC), with the Integrated Completed Likelihood (ICL) used to evaluate cluster separation. The resulting nine clusters are ordered by their means, and genes belonging to the highest-mean (two rightmost clusters with 29 and 147 genes) components correspond to those with the strongest contributions to the predictive models. The minimum normSHAP value within the selected clusters was used as an alternative threshold (th = 0.0107).

##### S6 - Associations between candidate genes and clinical phenotypes

To evaluate whether certain genes are more informative for specific demographic or clinical characteristics we performed an enrichment analysis of the candidate genes in clinical variables across all COPD samples using statistical tests appropriate to each variable type (Wilcoxon rank-sum [17], Student's t-test [18], or Pearson's chi-squared test [19]). Variables with adjusted p-values $\leq 0.05$ were considered statistically significant. These analyses revealed associations for several lung function measures, including emphysema, DLCO, and FEV1, while no enrichment was found for smoking status (Fig. 3c, Table S5).

The associated genes, all from the *extended* candidate set, reveal a multifaceted landscape of COPD pathology. These include markers linked to alveolar epithelial integrity (*SFTPC*), vascular remodelling and angiogenesis (*VEGFA*), and fibroblast activation (*PDGFRA*), as well as AGT, previously implicated in vascular dysregulation in COPD patients with comorbid hypertension [20–22]. Moreover, the results highlight cellular maintenance and immune disruption, through the enrichment of *TMEM138*, a cilia-related gene previously implicated in COPD deregulation, and *S100A10*, implicated in macrophage function and inflammatory responses [23]. Additional genes *CD164*, *CORO1C*, and *CAST* are involved in cell migration, cytoskeletal dynamics, and proteolytic regulation. Finally, other enriched genes such as *GAS2* and *GLIS1* emerged as DLCO-associated despite lacking well-established links to lung biology or COPD, suggesting potentially underexplored mechanisms contributing to disease heterogeneity.

It is important to note that, although no enrichment was observed for smoking status, several candidate genes (e.g., *CYP1B1*, *AZGP1*) are known to be associated with tobacco exposure [24, 25]. The lack of statistical signal likely reflects limitations in our clinical data, such as the absence of detailed smoking variables (e.g., packs/year).

Other highlighted candidate genes—including *MMP7*, *AHR*, *TGFB2*, and *STAT3*—were not significantly enriched in individual clinical variables, yet can be of interest to the disease. *MMP7*, while less studied than other matrix metalloproteinases (e.g. *MMP1*, *MMP2*, *MMP8*, *MMP9*), may reflect altered fibroblasts implicated in extracellular matrix remodelling mechanisms with higher variability. *AHR*, a regulator of xenobiotic response pathways, aligns with tobacco-related molecular stress, whereas nearby variants of *TGFB2* have been previously associated with emphysema. *STAT3*, a central signalling hub, may reflect broader inflammatory or regenerative programs, and association with other morbidities.

##### S7 - Clustering and subgroup clinical enrichment

To characterise patient-level heterogeneity, we performed hierarchical clustering using either aggregated normSHAP or expression levels of candidate genes with Euclidean distance. Clustering based on the aggregated normSHAP values per sample clearly separated disease from control samples, whereas separation was more challenging using traditional approaches (Fig. S21). Consistently, unsupervised clustering based only on the expression of the candidate genes stratified patients into COPD and control patients (Fig. S22).

Clinical enrichment analysis was subsequently performed within each identified subgroup (of the expression-based clustering) to assess subgroup-specific molecular and phenotypic patterns. Cluster 2 comprised control samples, representing healthier phenotypes and characterised by non-smokers (p-value $= 7.496 \times 10^{3}$), better lung function and the absence of COPD-related features. Cluster 3 delineated a severe COPD expression endotype (p-value $= 3.998 \times 10^{-3}$), associated with advanced airflow limitation (predicted FEV1 $= 1.516 \times 10^{-4}$, predicted FVC $= 4.013 \times 10^{-3}$) and increased emphysema (p-value $= 1.129 \times 10^{-4}$). Cluster 5 captured a more heterogeneous, core COPD transcriptional signature, enriched in smoking (current and ever categories, p-value $= 1.499 \times 10^{-2}$) and associated with reduced lung function (predicted FEV1 $= 7.2569 \times 10^{-5}$, predicted FVC $= 8.639 \times 10^{-3}$), emphysema and DLCO (p-value $=$ $1.625 \times 10^{-4}$, p-value $= 1.168 \times 10^{-7}$). Finally, Cluster 1 did not show significant enrichment for specific phenotypic variables, likely reflecting the intrinsic molecular heterogeneity of COPD and the coexistence of multiple disease mechanisms within this group (Table S4).

##### S8 - Cross-tissue expression analysis

Tissue-level expression data were retrieved from the GTEx Portal (GTEx Analysis V10 release accessed 12/2025) to assess the expression patterns of candidate genes across human tissues, with particular emphasis on lung enrichment and multi-organ expression profiles.

*Canonical* candidate genes, identified through classical approaches, tend to show lung-enriched expression, consistent with their established roles in COPD pathogenesis (Fig. S17). In contrast, *extended* candidate display a more diverse expression profile (Fig. S18). While several *extended* genes were also lung-enriched (e.g., *SFTPC*, *TNNC1*, *S100A8*), many were expressed across multiple tissues, suggesting potential systemic involvement.

Notably, a subset of both *canonical* and *extended* genes show blood expression together with lung (e.g. *S1AA08*, *TNFRSF1B*, *PI4KA*, *S100A10*, *MAPK14*, *CHST15*, *MMP9*, *PROK2*, *FCN1*, *CTSG*, *OSM*, *CLC*, *CHI3L1*), indicating potential as non-invasive biomarkers. Certain candidates exhibit enriched expression in lung and other organs, including *BTNL9*, *AZGP1* (lung and breast), *CYP1B1* (lung, uterus and fallopian), *PDGFRA* (lung and ovary), or genes expressed in lung and liver/brain, which may reflect sex-specific effects [26], comorbidities (Fig. S19)[27], or environmental exposures such as smoking [24, 25].

To further evaluate cellular specificity within the lung, expression of *extended* candidate genes was examined using Human Cell Atlas (HCA) lung data, confirming their representation across relevant pulmonary cell types (Fig. S20) [28].

##### S9 - Shared molecular signatures across chronic lung diseases

Our final candidate list of genes included both well-known COPD-associated genes (e.g. *CYTH1*, *COX7B*, *ZDHHC11*, and *C1RL*) [29–32] and novel potential biomarkers. Notably, some of the identified genes have been previously implicated in biological processes shared with other chronic respiratory diseases, such as asthma (e.g. *KIAA0284*, *ITLN1*) [33]. In addition, several candidates have been reported in conditions that frequently co-occur with COPD or share common risk factors, including lung cancer (e.g. *ASCL1*, *ROR1*)[34, 35] and idiopathic pulmonary fibrosis (e.g. *LRRFIP1*, *AGER*) [36].

These associations might arise from inflammation-driven alterations in epithelial populations (e.g. AT1 and AT2 cells), but also from transcriptional changes in COPD-associated fibroblasts, including genes such as *VEGFA*, *CCL19*, *SFTPC* and *CR2* [7].

#### 2. SUPPLEMENTARY FIGURES

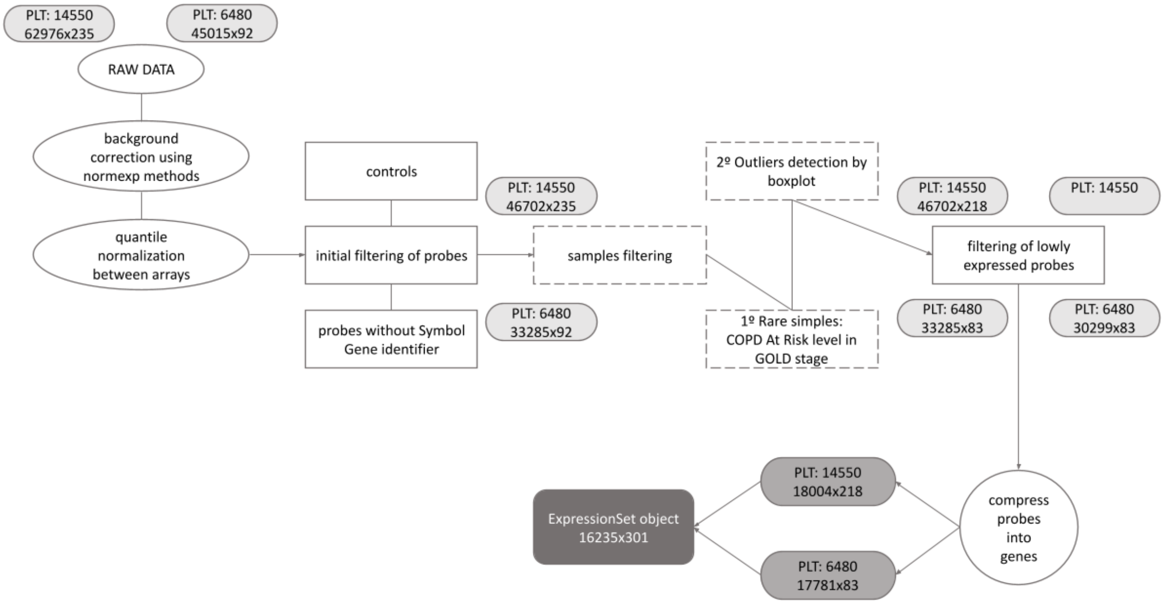


**Fig. S1. Pipeline for processing microarray data.** Collection of preprocessing steps applied to the two Agilent microarray platforms (PLT:14550 and PLT:6480) separately until their union. Ovals represent the first pre-processing steps that should be done over the microarray data (background correction using normexp method and quantile normalisation between arrays). Rectangles show the filtering steps of the probes (continuous) and samples (dotted). First, filtering of control probes and those that have no correspondence with any GeneSymbol identifier (platform annotations were downloaded from GEO). After that, samples wrongly annotated (that is, COPD patients marked as being controls in the GOLD stage category) were deleted. Moreover, we used the Kolmogorov-Smirnov statistic to compare each array's intensity distribution and the distribution of the pooled data for obtaining the outlier samples. As this method bases its results on a simulated p-value, we generated 10000 randomizations and selected as outliers those samples that appear in at least 25% of the trials. Then, lowly expressed probes were also filtered, that is, probes with an expression count lower than half of the samples in the disease condition, with fewer samples ($>$ 42 in PLT:14550 and $>$ 8 in PLT:6480). Finally, we compressed the probes into genes to obtain our final expression object. Note that light grey figures show the dimensions of each platform in terms of probes, the grey ones in terms of genes, and the dark grey figure shows the dimensions of the final expression set object (once both platforms were joined).

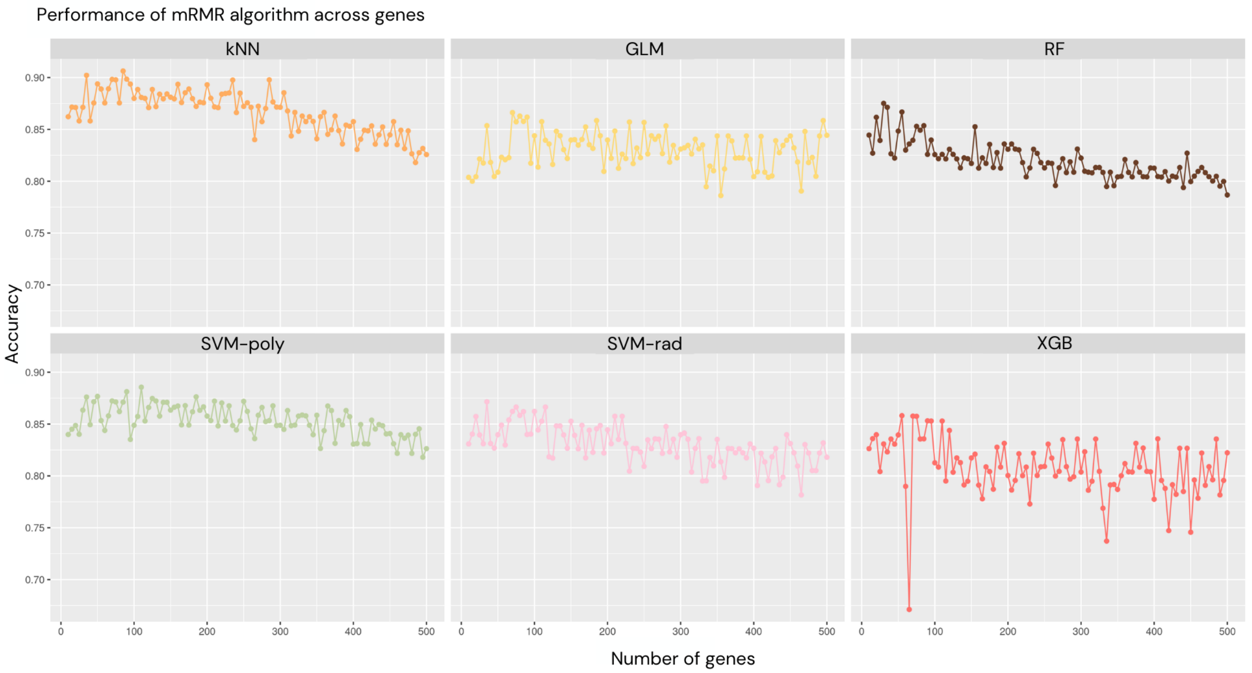


**Fig. S2. mRMR accuracy performance across genes.** These curves show how the prediction performance of the different classifiers (represented with different colours in the plot) built, including as input the top $k$ gene in the mRMR list, with $k \in [10, 500]$ behaves. The curves are based on the cross-validation accuracy achieved on the Bayes optimisation tuning methodology.

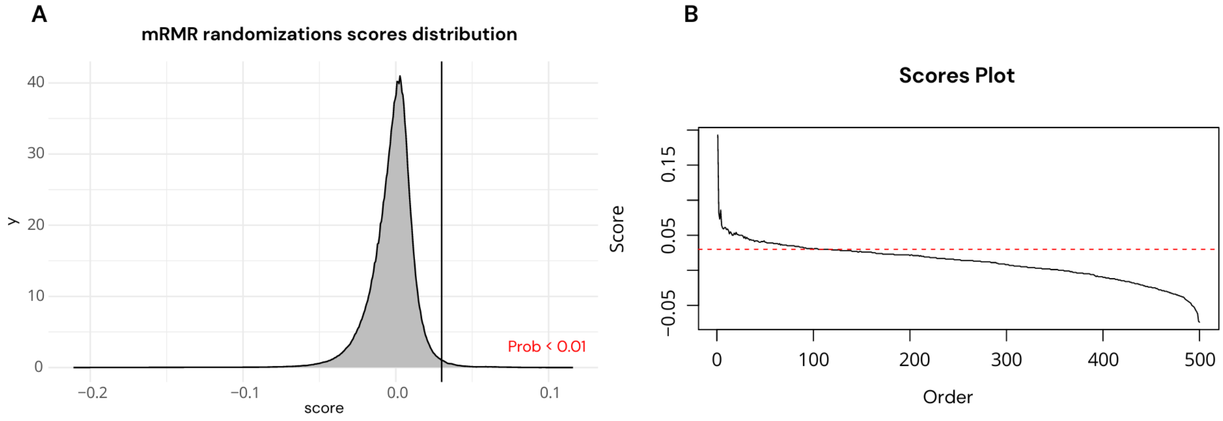


**Fig. S3. mRMR randomization scores distribution and top 500 real scores.** (A) shows the mRMR scores distribution obtained by running the mRMR algorithm on 1000 datasets with sample disease labels altered. The vertical line represents the threshold, $a$, for which the probability of a random variable, $x$, falling in the interval $[a, 1]$ is equal or less than 0.01. (B) shows the curve of the scores for the top 500 features selected by the algorithm when applied to the real training data. The horizontal line indicates the threshold $a = 0.03$ used to identify a gene as significant for distinguishing between COPD and control samples.

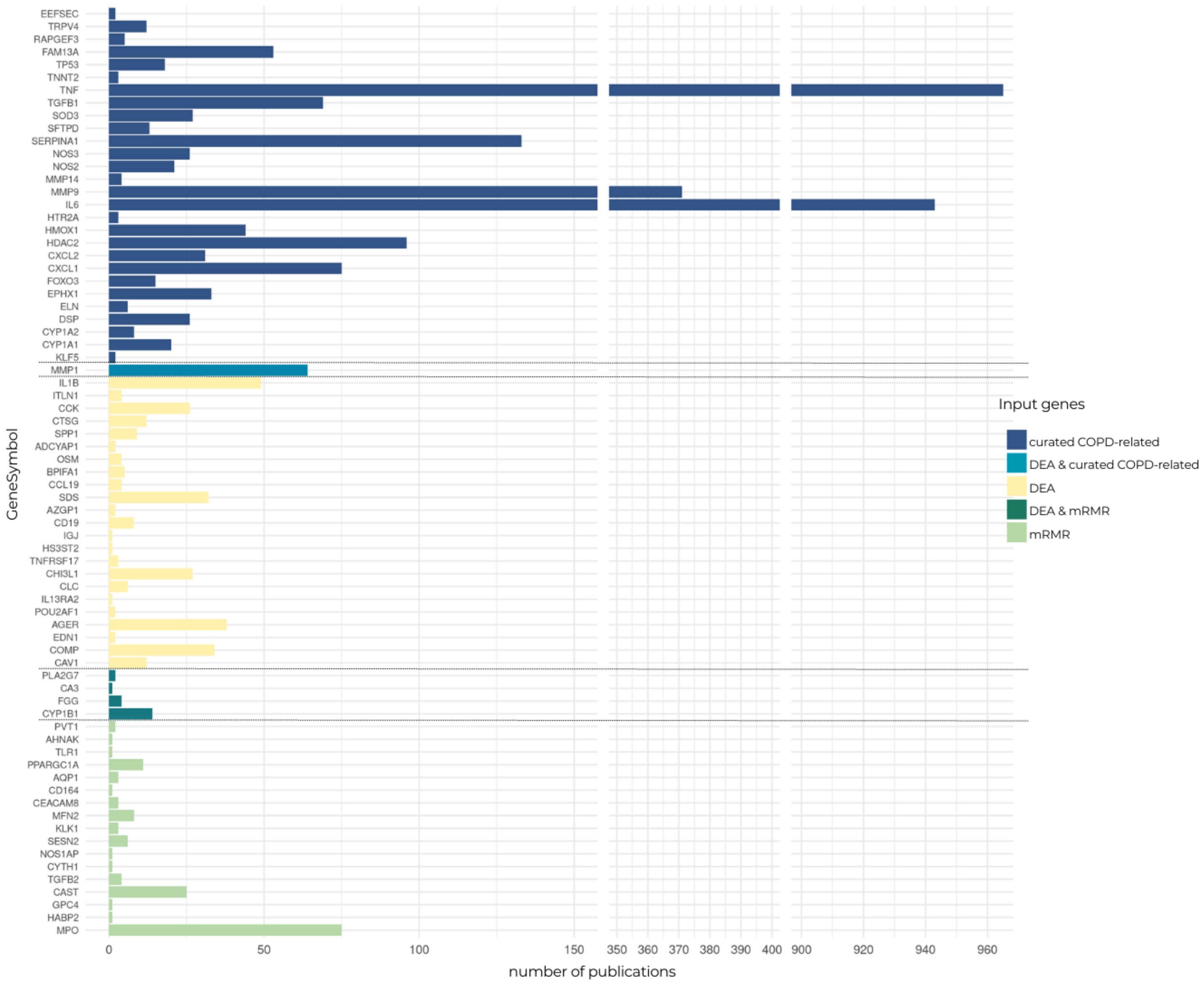


**Fig. S4. Literature-based validation of seed gene sets associated with COPD.** Confirmation of the association between the seed gene sets and COPD through a literature review using the PubMed database with the query defined in Note S3. The figure shows the genes with publications co-mentioned with COPD, along with the corresponding number of publications per gene. Colours represent the seed gene sets: COPD-related curated, DEA and mRMR, as well as their intersections (COPD-related curated∪DEA, DEA∪mRMR). Dashed horizontal lines separate the different gene groups for clarity.

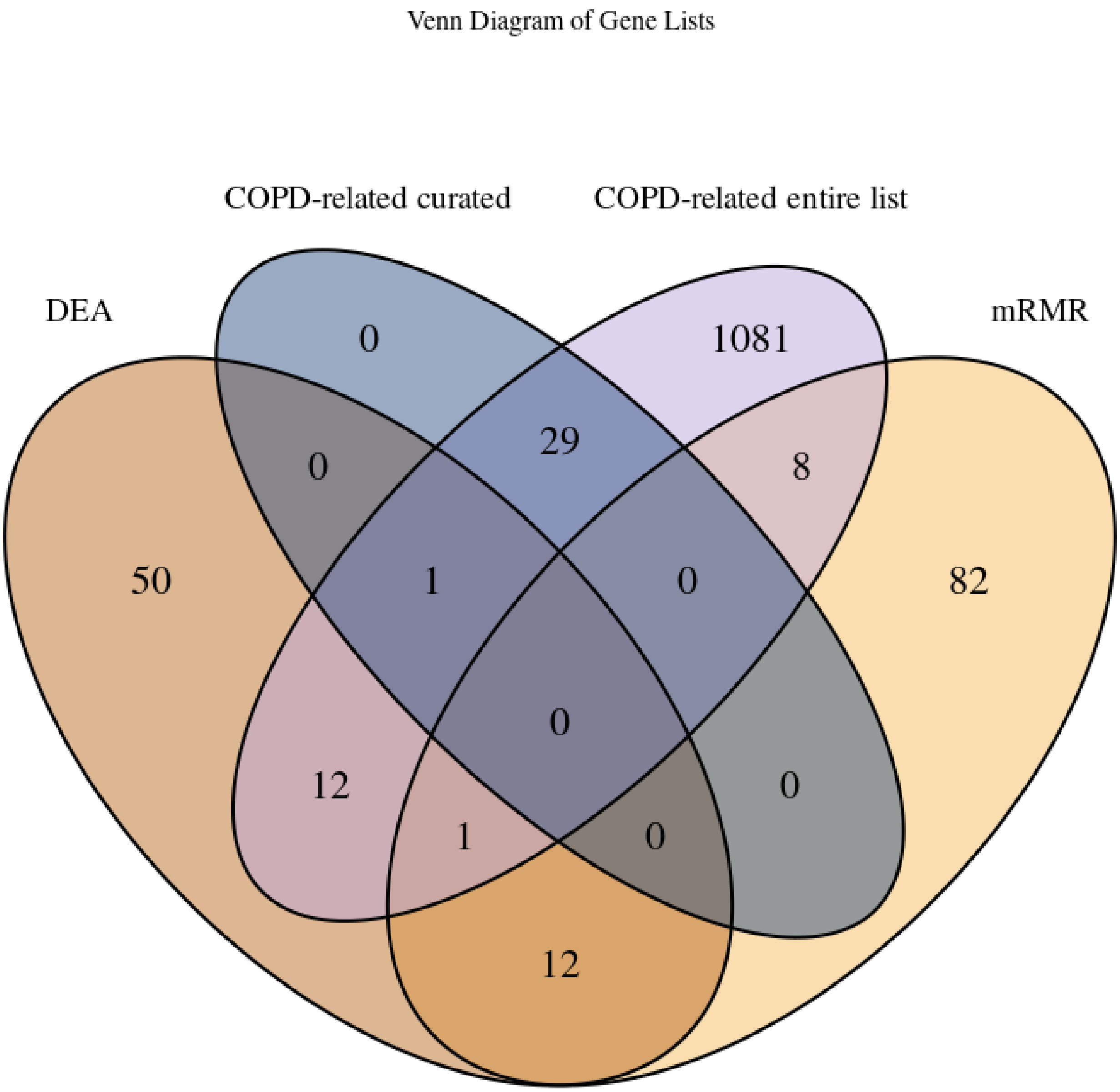


**Fig. S5. Intersection among seed gene lists used for feature selection.** Venn Diagram showing the overlap among the initial seed gene sets used in the analysis. Data-Driven (resulting from combining DEA genes and mRMR genes) and COPD-related genes (both curated and entire list) retrieved from DisGeNET.

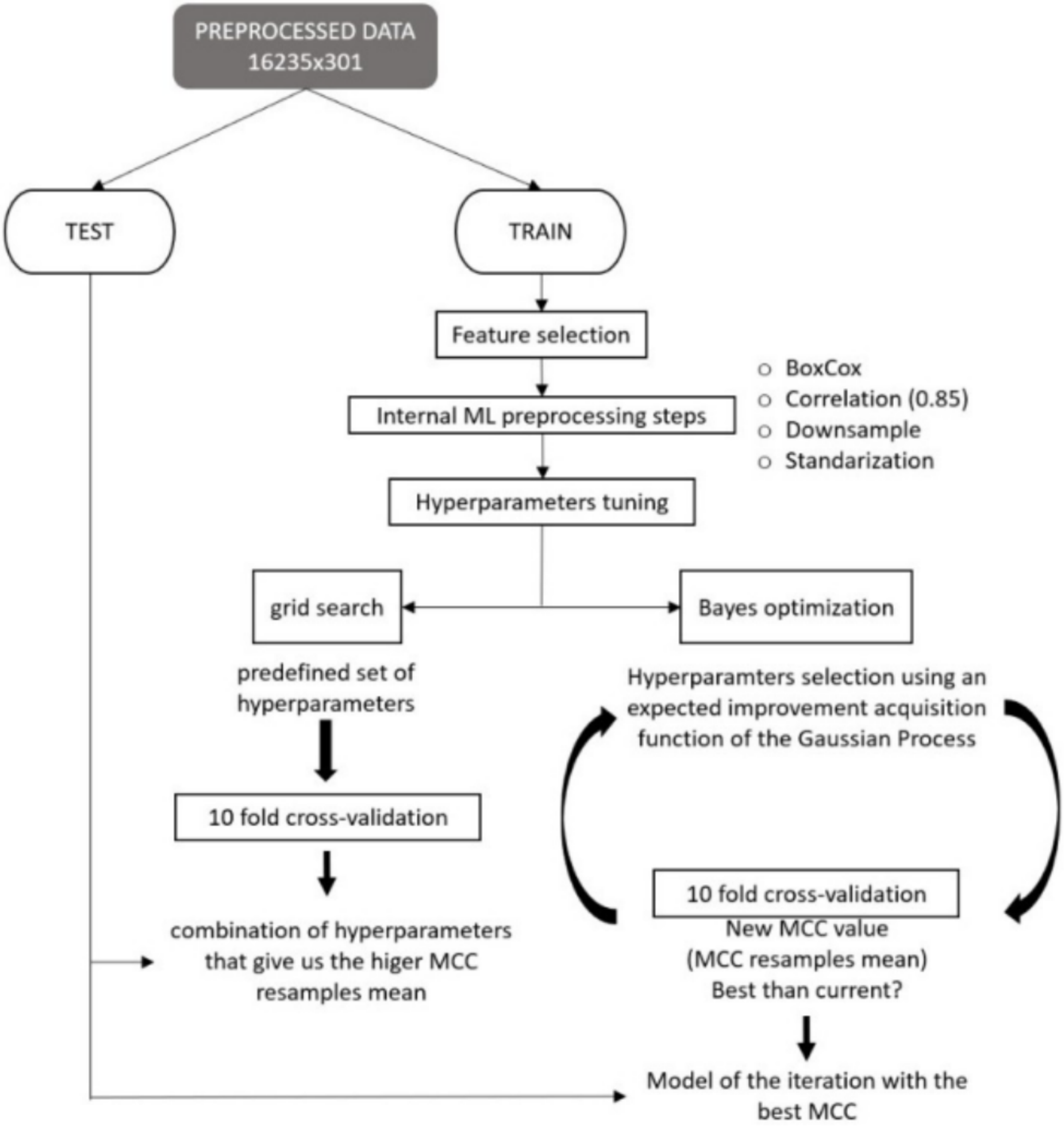


**Fig. S6. Training Procedure** After data splitting and feature selection, a standardised preprocessing recipe was applied to the training set, following the requirements of each classifier. Consequently, two hyperparameter tuning methodologies were evaluated: grid search and Bayesian optimisation, with the latter selected for the final models. Classifiers were trained on the complete training dataset, internally validated using 10-fold cross-validation, and subsequently evaluated on the independent test set.

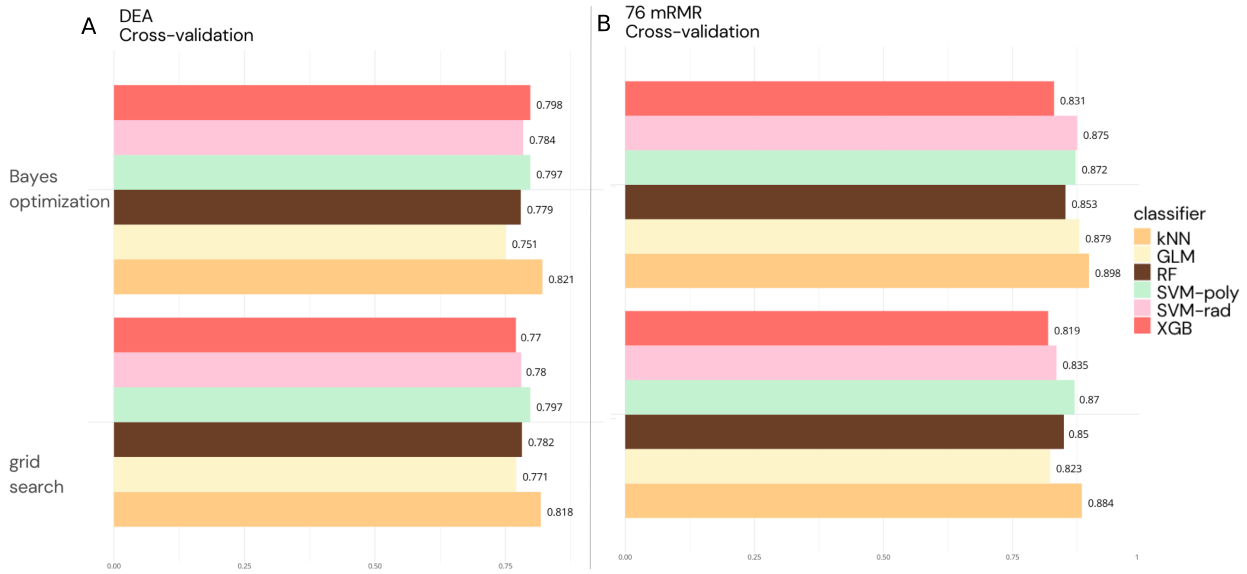


**Fig. S7. Tuning methodologies comparison.** Accuracy comparison between two tuning methodologies: (A)Bayes optimisation (top) and grid search (bottom). Cross-validation performance of DEA (A) and 76 mRMR genes (B) are shown.

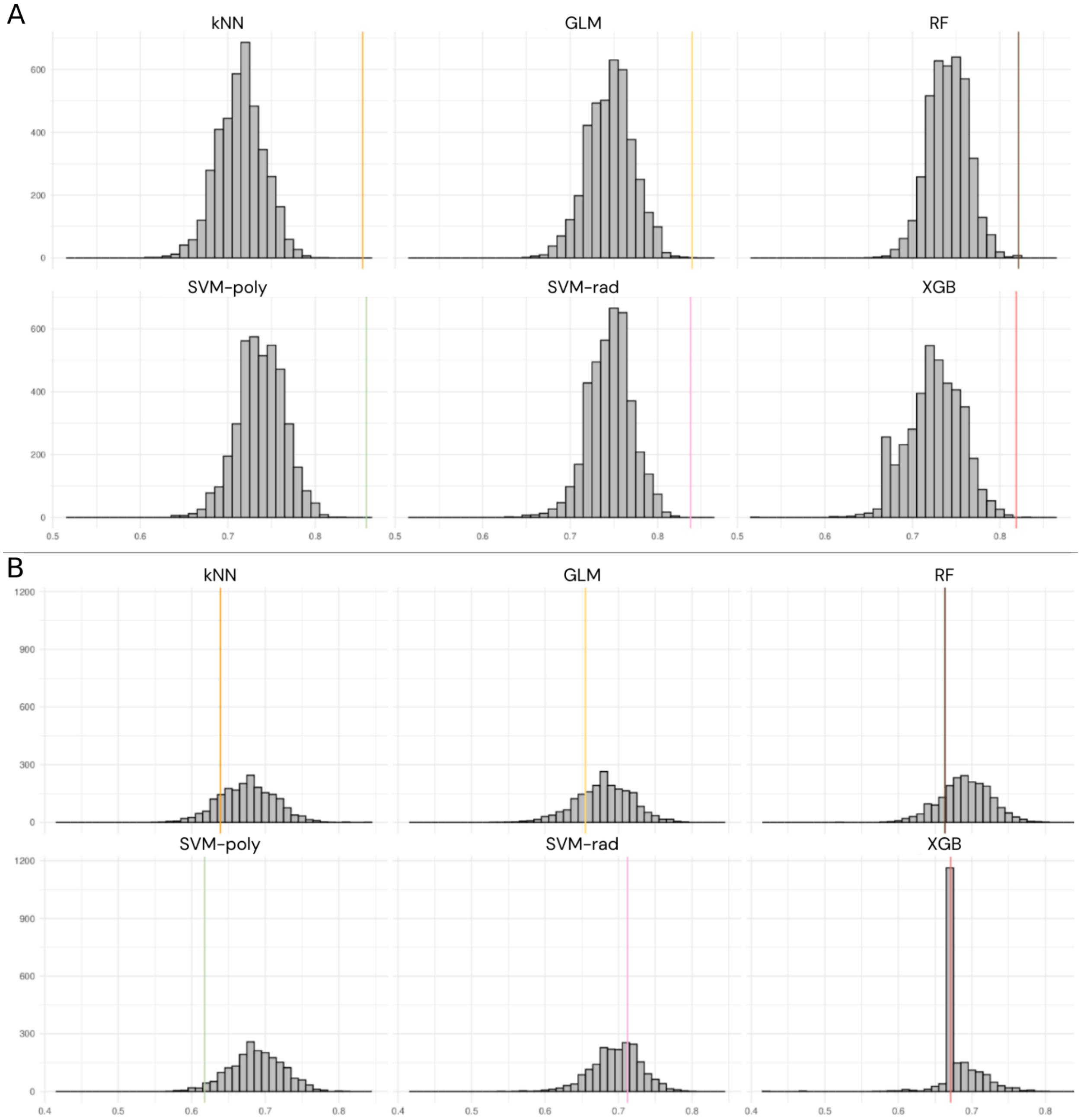


**Fig. S8. Comparison of model performance using random gene sets versus selected seed gene sets.** (A) Distribution of classification accuracies obtained from 1,000 models trained using 166 randomly selected genes as input features. (B) Distribution of classification accuracies obtained from 1,000 models trained using 30 randomly selected genes as input features. In both plots, vertical coloured lines indicate the accuracy achieved using the predefined gene sets: 166 data-driven genes (A) and 30 COPD-related curated genes (B). These comparisons assess whether the selected gene sets outperform random expectations.

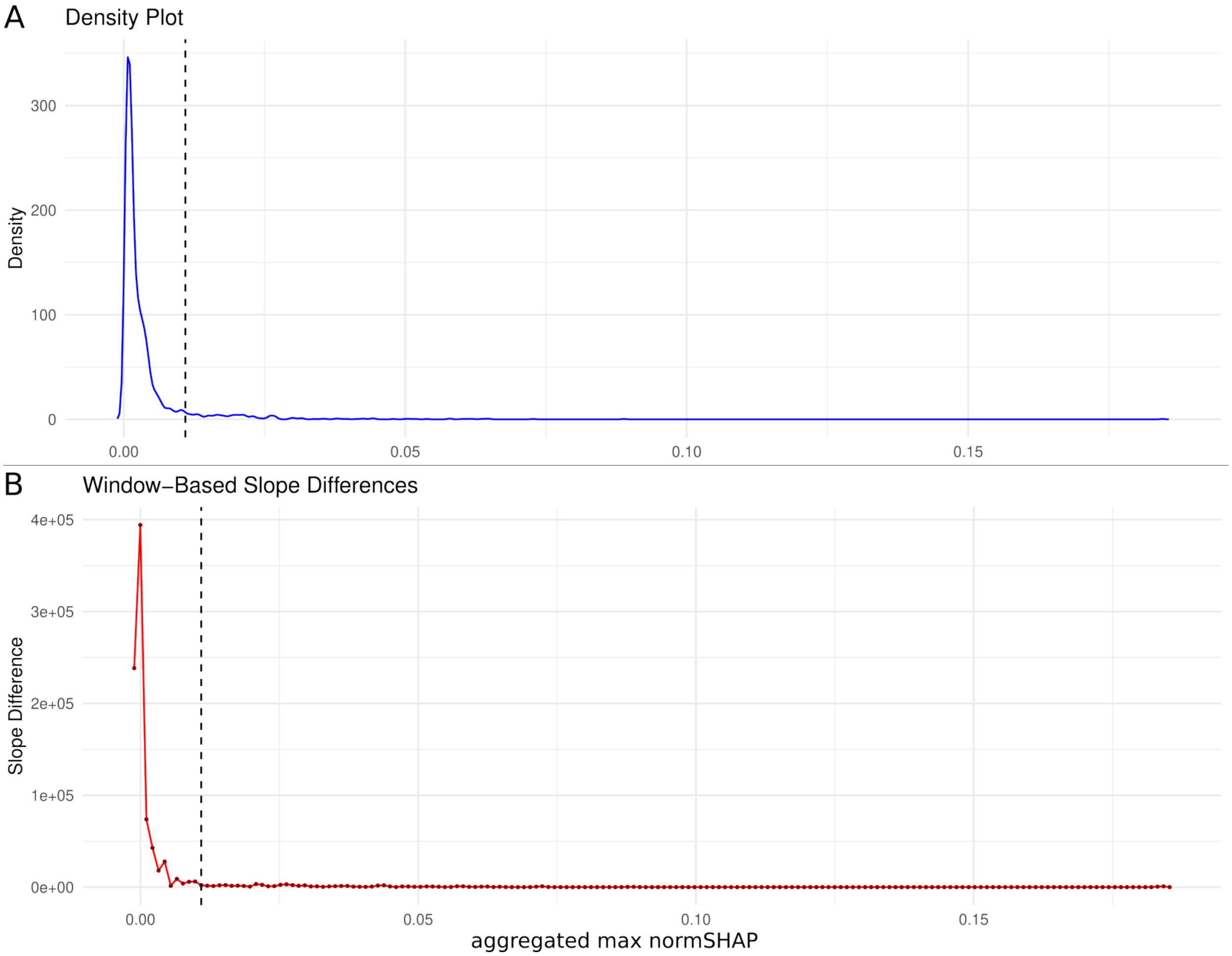


**Fig. S9. Candidate gene selection by using a density-based approach.** (A) Density plot of aggregated normSHAP values analysed using a window-based approach. (B) displays the corresponding differences in density and slope across the range of aggregated normSHAP values, supporting the selection of the stabilisation threshold. The dashed vertical line indicates the threshold (th = 0.0109) at which the density curve stabilises, identified by tracking changes in density and slope within fixed-size windows (0.001).

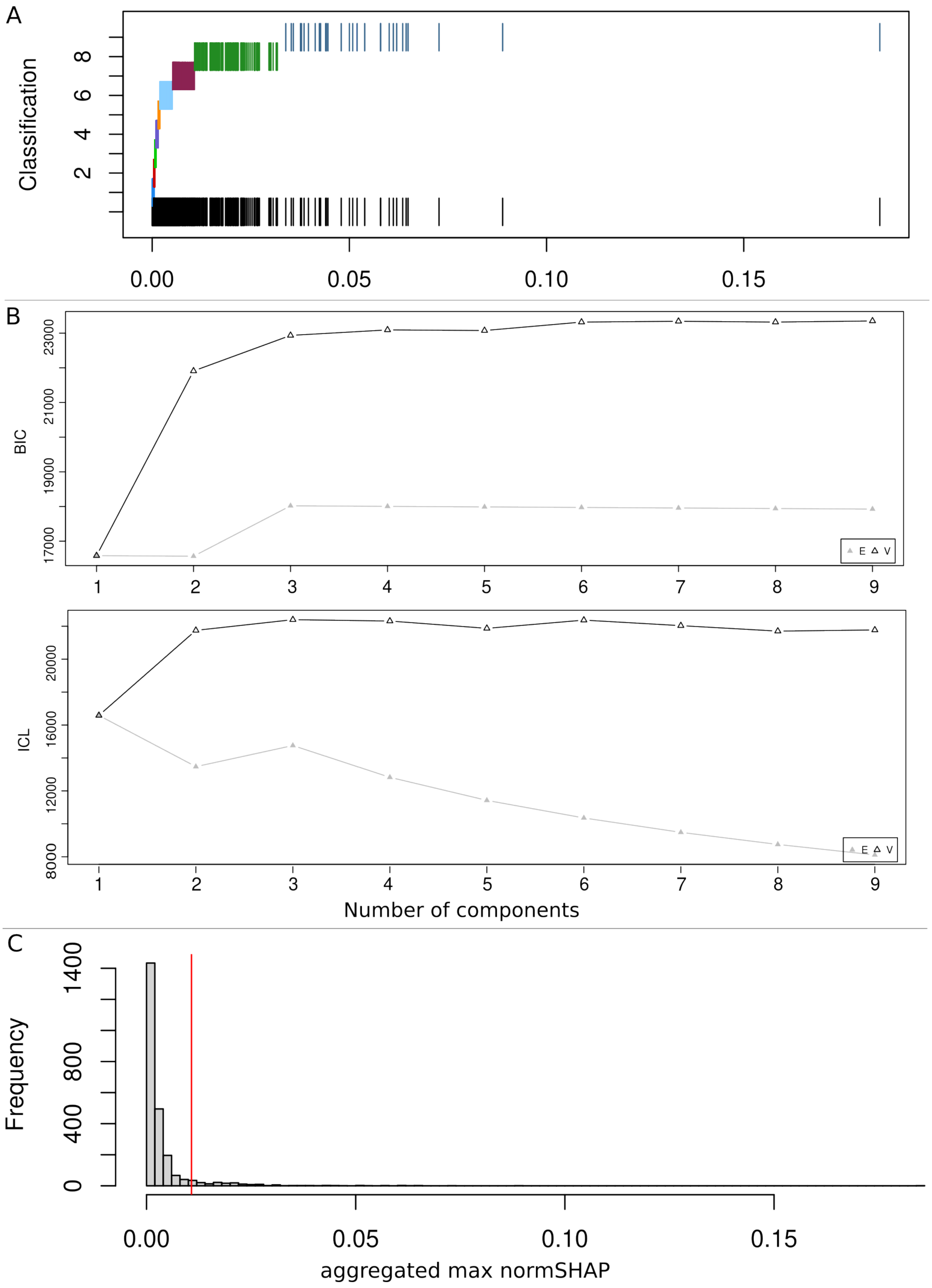


**Fig. S10. Results of Gaussian mixture model clustering applied to SHAP values.** The top plot displays the classification of SHAP values into clusters. The bottom histogram shows the SHAP value distribution, with a red vertical line marking the threshold derived from the minimum value of the selected clusters (th = 0.0107). This threshold identifies genes with the highest contribution to the model.

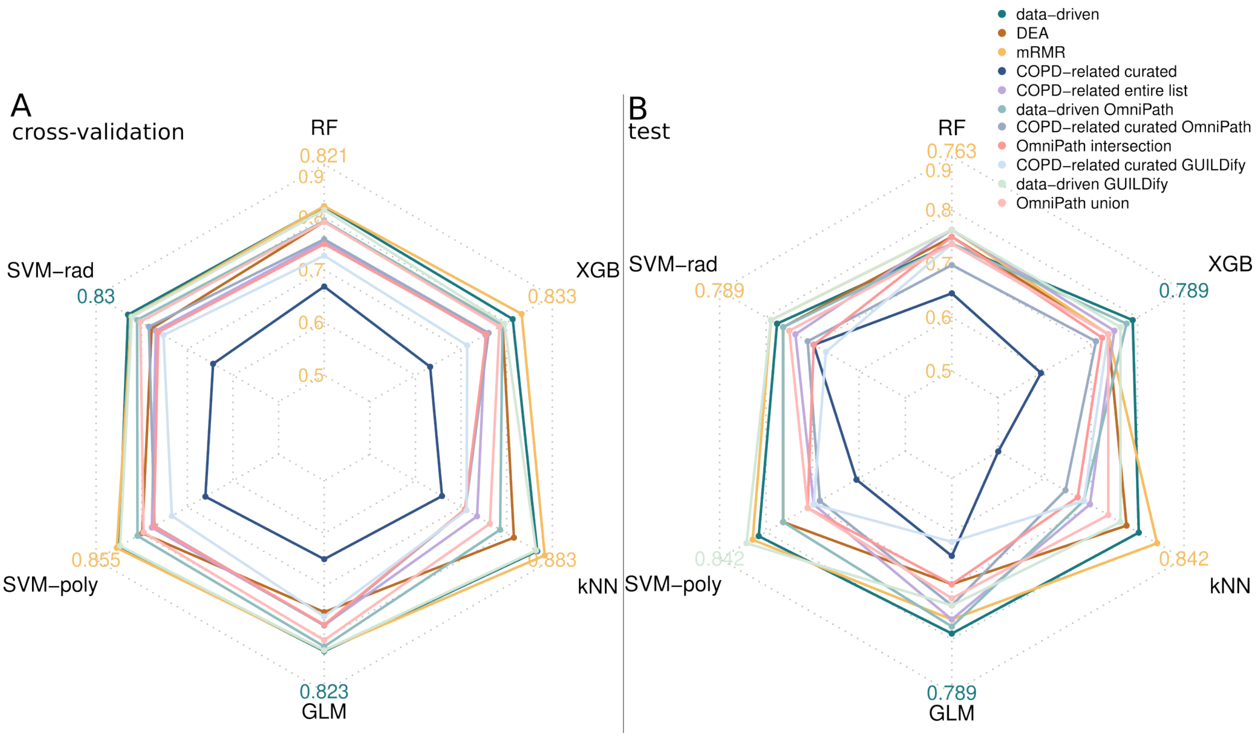


**Fig. S11. Prediction performance across all scenarios.** Accuracy scores for each ML model (vertices), with colours indicating the input gene set. A panel shows cross-validation results; B panel shows test results.

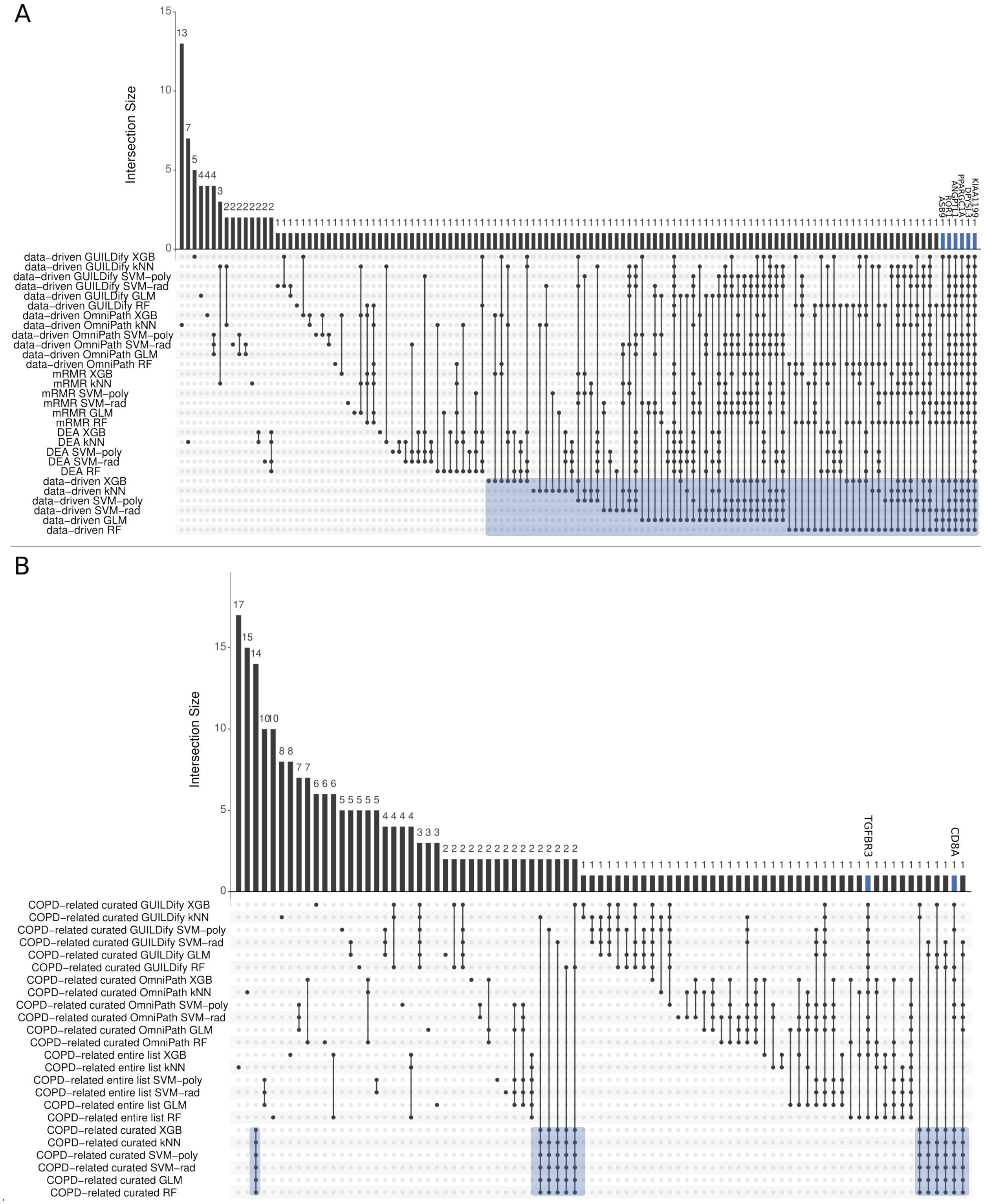


**Fig. S12. Overlap of top 30-ranked genes across scenarios.** (A) Overlap of the top 30 genes ranked by normSHAP values across classifiers for data-driven gene sets (DEA, mRMR, data-driven) and their expansions using OmniPath and GUILDify. Genes highlighted in blue appear in more than two-thirds of the 30 total scenarios (*KIAA1199*, *DPYSL3*, *ROR1*, *PPARGC1A*, *ANGPTL1*, and *ASB9*). (B) Overlap of the top 30 normSHAP-ranked genes across classifiers for COPD-related gene sets (curated from DisGeNET) and their expansions. Genes highlighted in blue appear in half of the combinations (*D8A* and *TGFBR3*). In both plots, blue squares represent the seed gene sets.

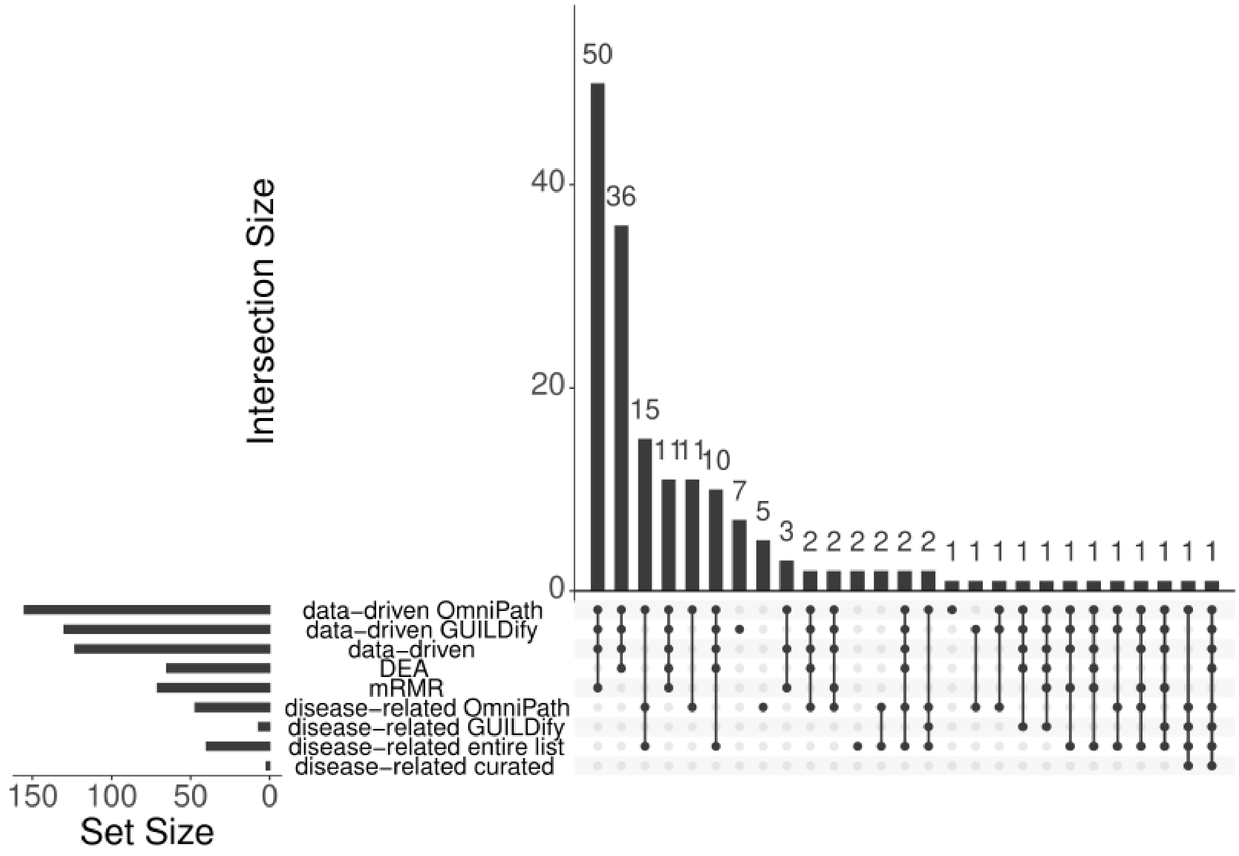


**Fig. S13. Origin of candidate genes across input gene sets.** UpSet plot summarising the overlap and unique contributions of candidate genes according to the gene sets used as input to the classifiers.

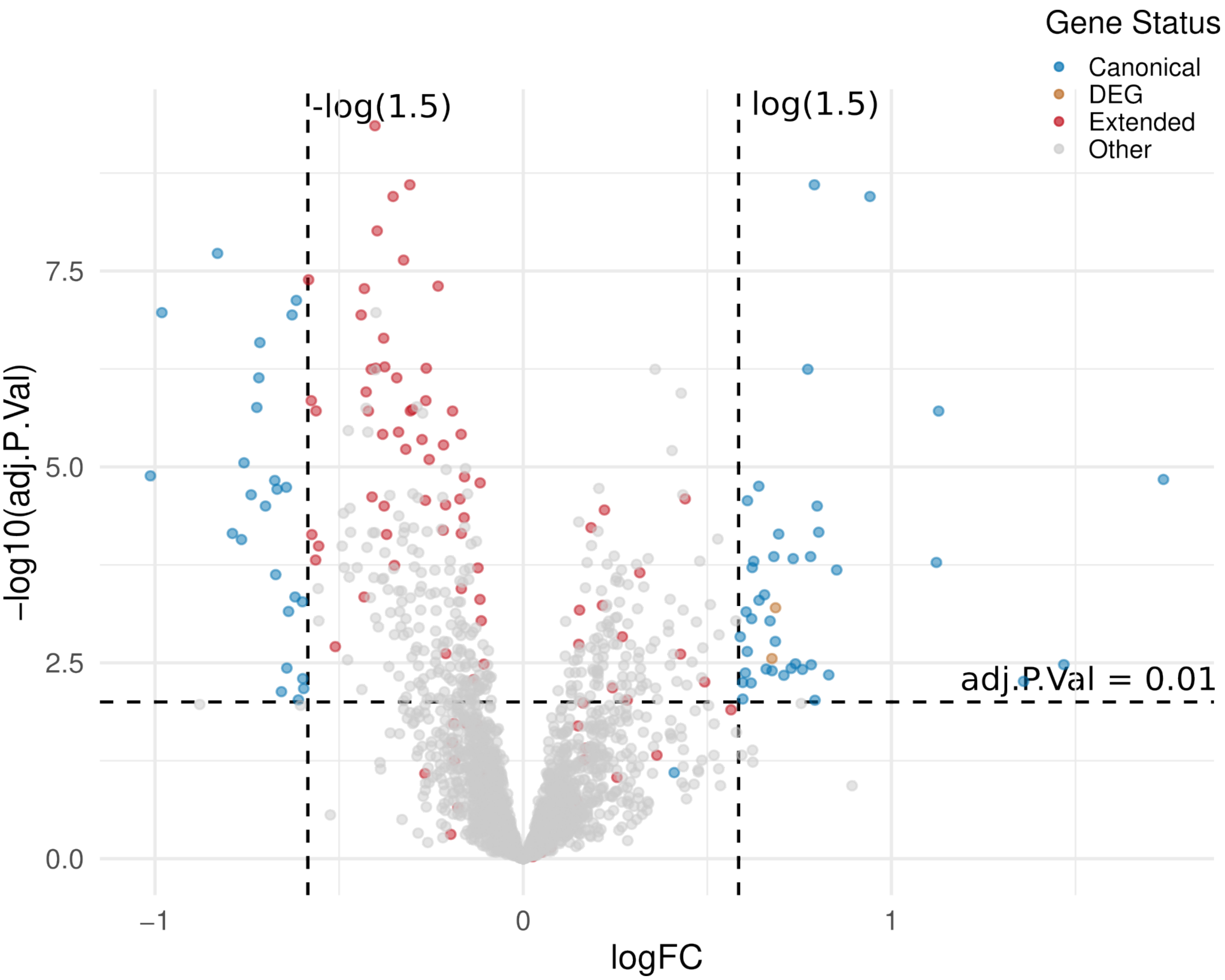


**Fig. S14. Candidate genes in DEA.** Volcano plot showing the position of candidate genes in the DEA results, relative to statistical significance (FDR $\leq$ 0.01) and effect size (FC $\geq$ 1.5). Candidate genes are highlighted to illustrate their distribution across significant and non-significant regions and coloured according to their classification: *canonical* (blue), *extended* (red), and genes identified exclusively by DEA but not included in the final candidate set.

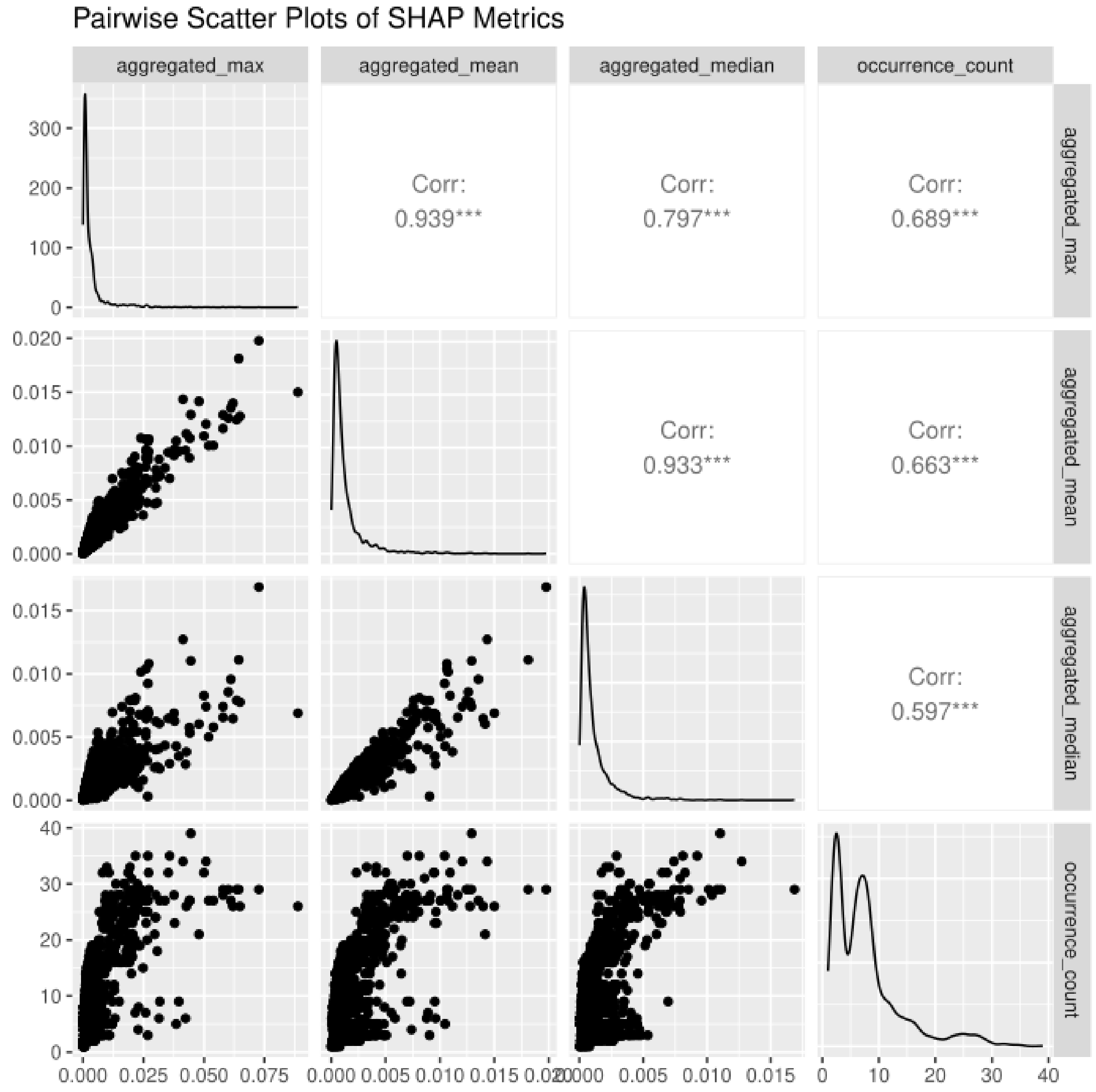


**Fig. S15. Pairwise Scatter Plots of SHAP metrics.** Scatter plot showing the pairwise Spearman correlation and the distribution of each aggregated SHAP metric (aggregated maximum, mean and median, and occurrence count).

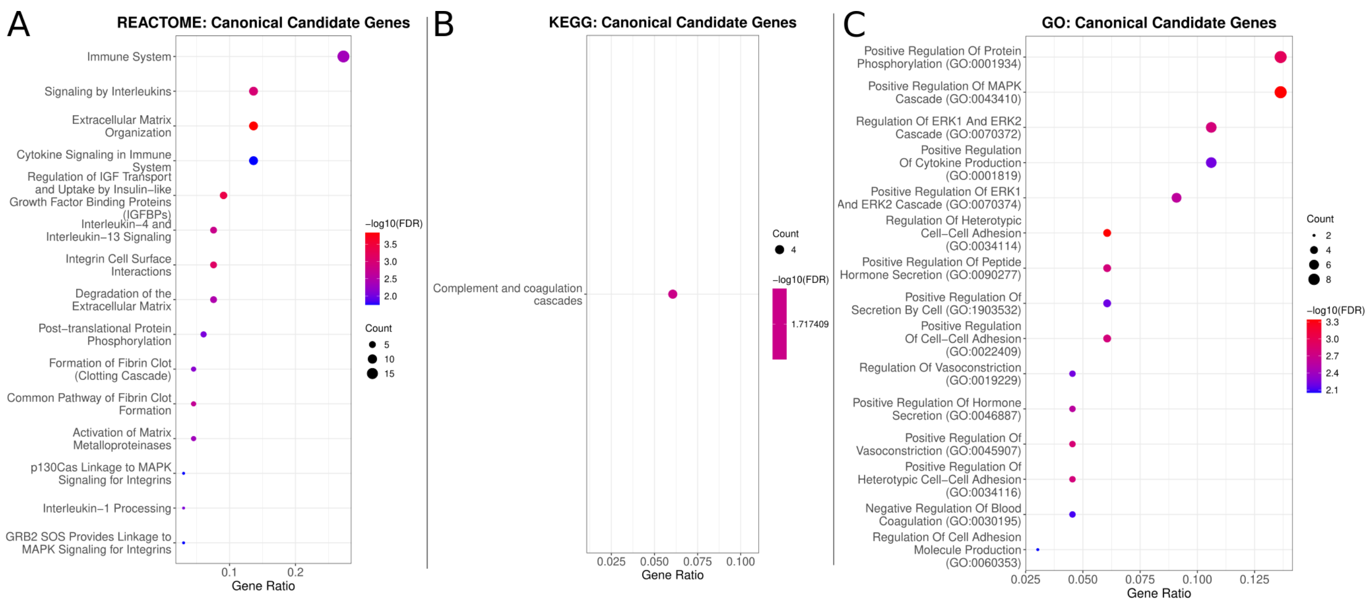


**Fig. S16. Functional enrichment analysis of *canonical* genes.** Top 20 enriched pathways for *canonical* genes identified using (A) Reactome, (B) KEGG, and (C) Gene Ontology (GO). In each panel, the x-axis represents the gene ratio, the y-axis lists enriched pathways, colour encodes the significance level as -log10(FDR), and point size corresponds to the number of genes contributing to each pathway (count).

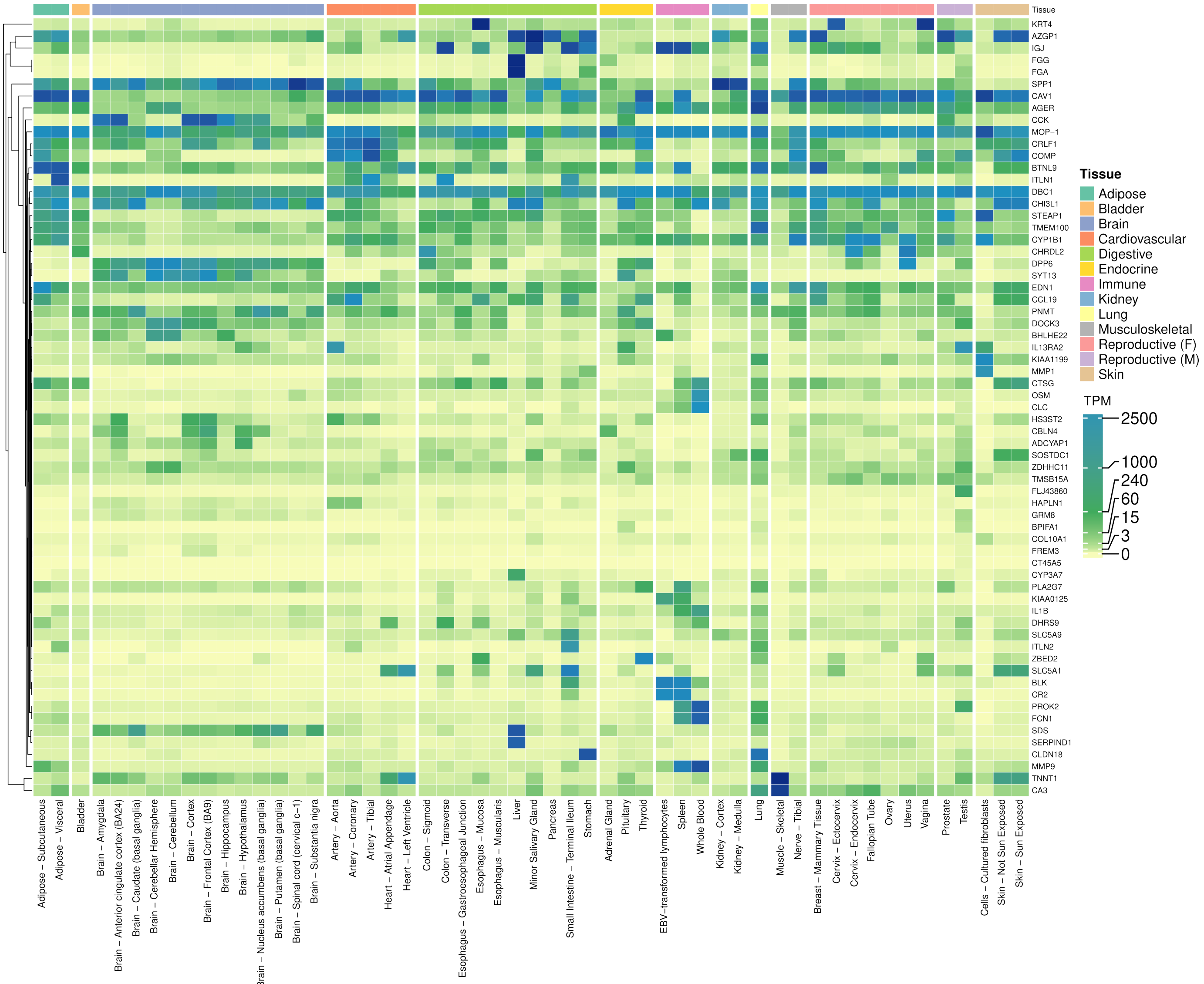


**Fig. S17. GTEx tissue expression heatmaps of *canonical* candidate genes.** Heatmaps showing tissue-specific expression profiles of *canonical* candidate genes across human tissues from the GTEx project. The x-axis represents tissues, and the y-axis represents genes. Colour intensity indicates expression levels measured as Transcripts Per Million (TPM). Top annotations group tissues into broader anatomical categories, facilitating comparison of tissue-specific patterns. The data used for the analysis described here were obtained from the GTEx Portal on 12/2025.

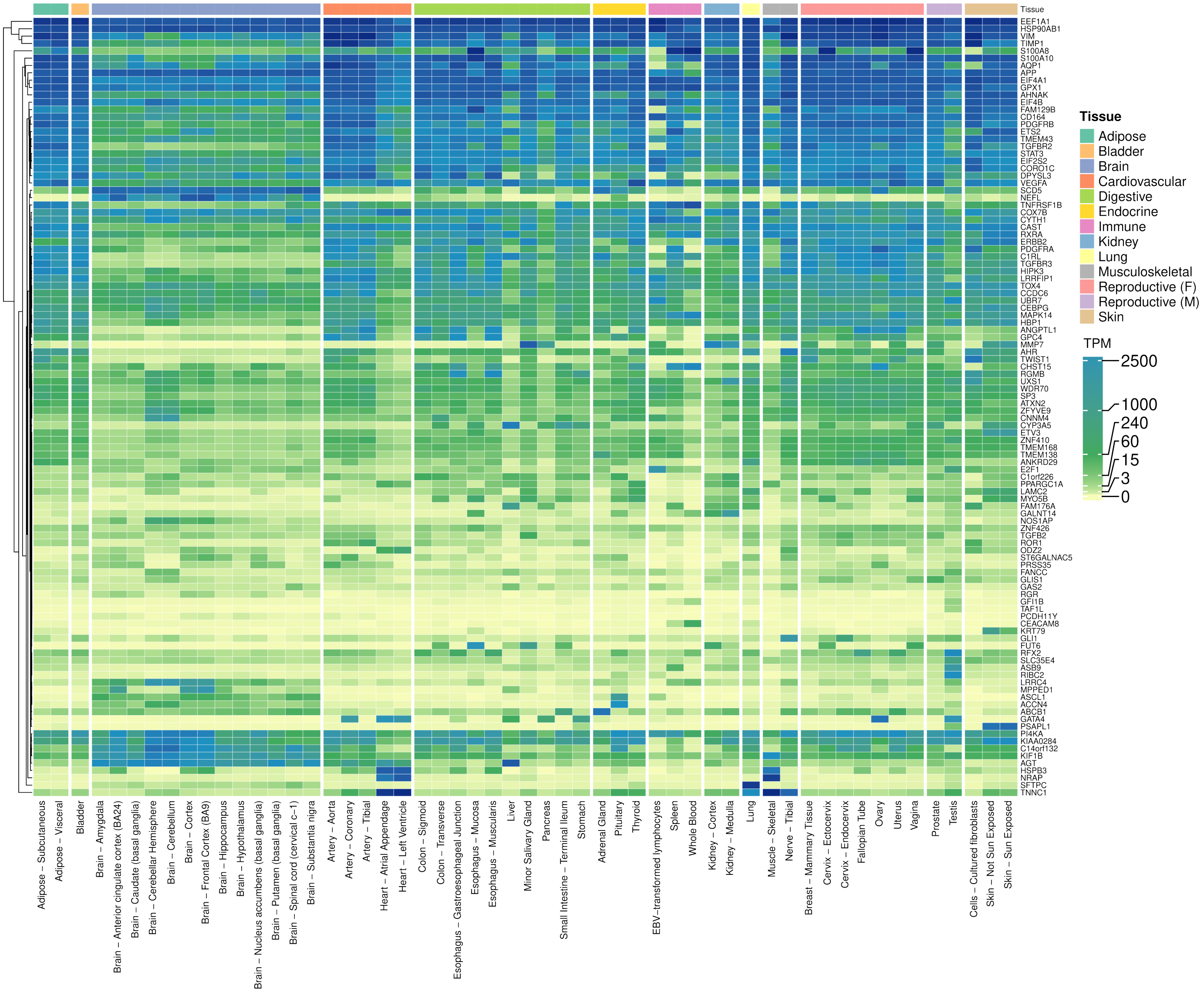


**Fig. S18. GTEx tissue expression heatmaps of *extended* candidate genes.** Heatmaps showing tissue-specific expression profiles of *extended* candidate genes across human tissues from the GTEx project. The x-axis represents tissues, and the y-axis represents genes. Colour intensity indicates expression levels measured as TPM. Top annotations group tissues into broader anatomical categories, facilitating comparison of tissue-specific patterns. The data used for the analysis described here were obtained from the GTEx Portal on 12/2025.

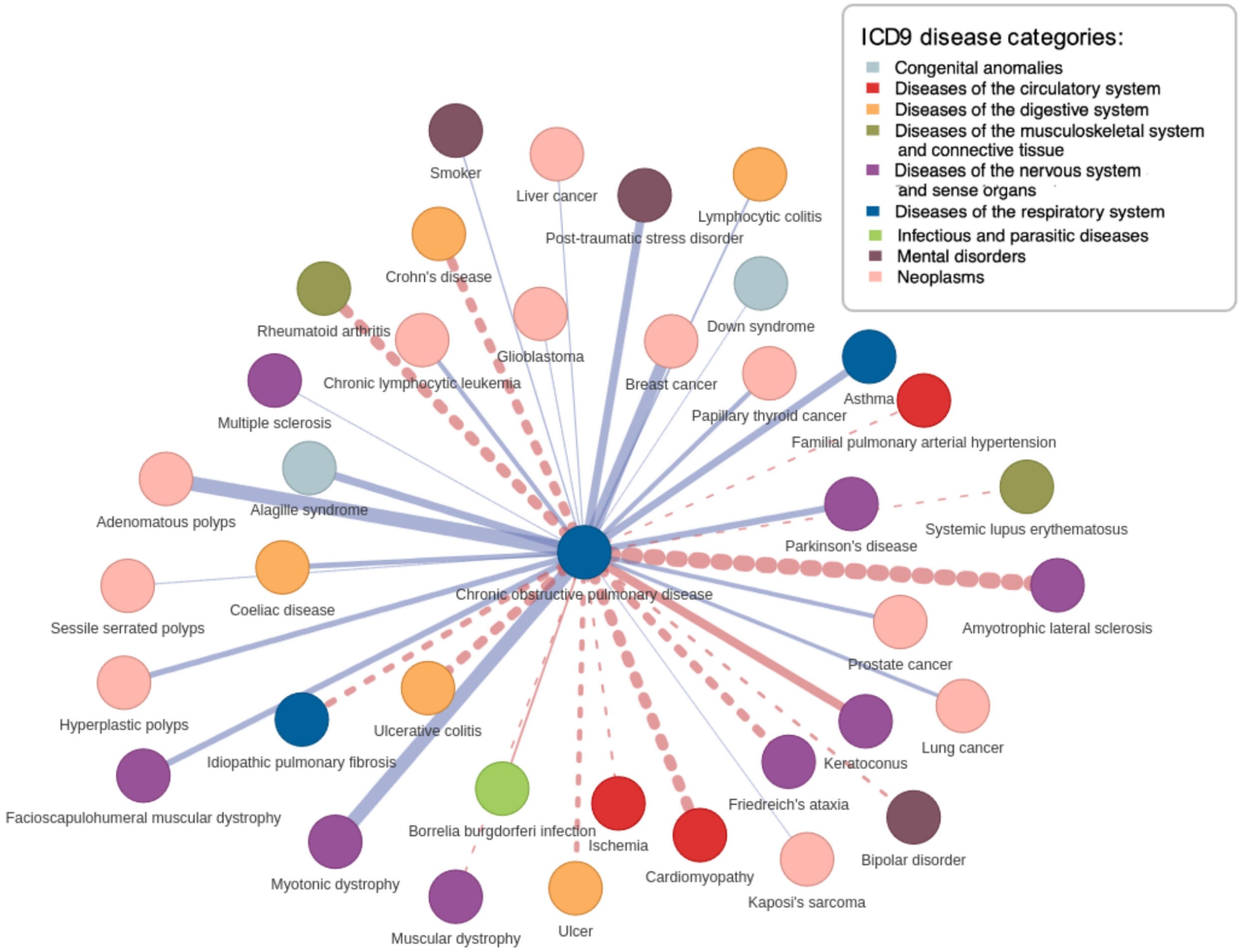


**Fig. S19. Disease similarity network based on transcriptomic profiles highlighting COPD comorbidities.** The network was generated using the Disease Perception platform (http://disease-perception.bsc.es/rgenexcom/. Nodes represent diseases, and edges indicate significant transcriptomic similarity between disease expression profiles. COPD is in the centre of the network. Red edges denote positive interactions and blue edges negative interactions. Node colours represent ICD9 disease categories.

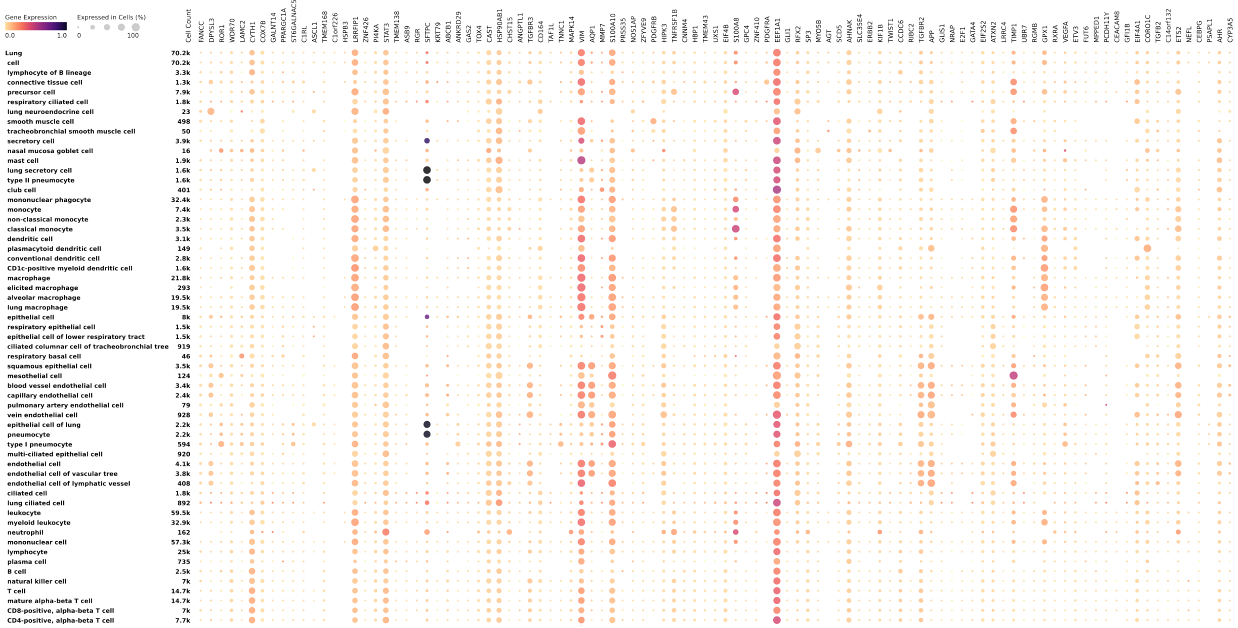


**Fig. S20. Cell-type enrichment of *extended* candidate genes in the human lung.** Dot plot showing the expression of *extended* candidate genes across lung cell types using data from the Human Lung Cell Atlas [28]. Dot size represents the proportion of cells expressing each gene within a given cell type, while colour intensity reflects the average expression level.

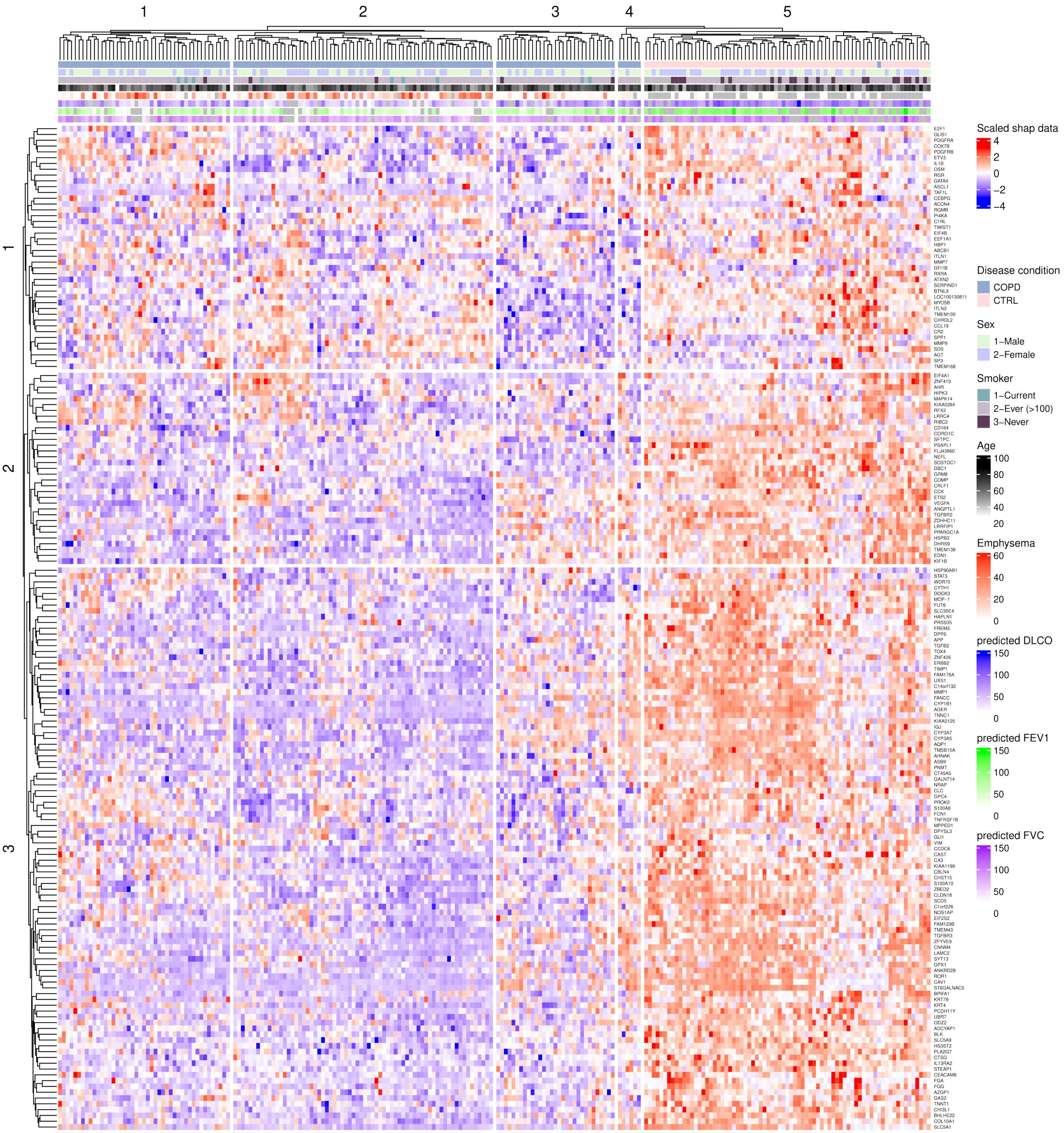


**Fig. S21. Patient clustering based on aggregated normSHAP values.** Heatmap showing hierarchical clustering of patients (columns) and candidate genes (rows) using aggregated normSHAP values per sample. Colour intensity reflects the scaled contribution of each gene to the model prediction. Top annotations display clinical characteristics.

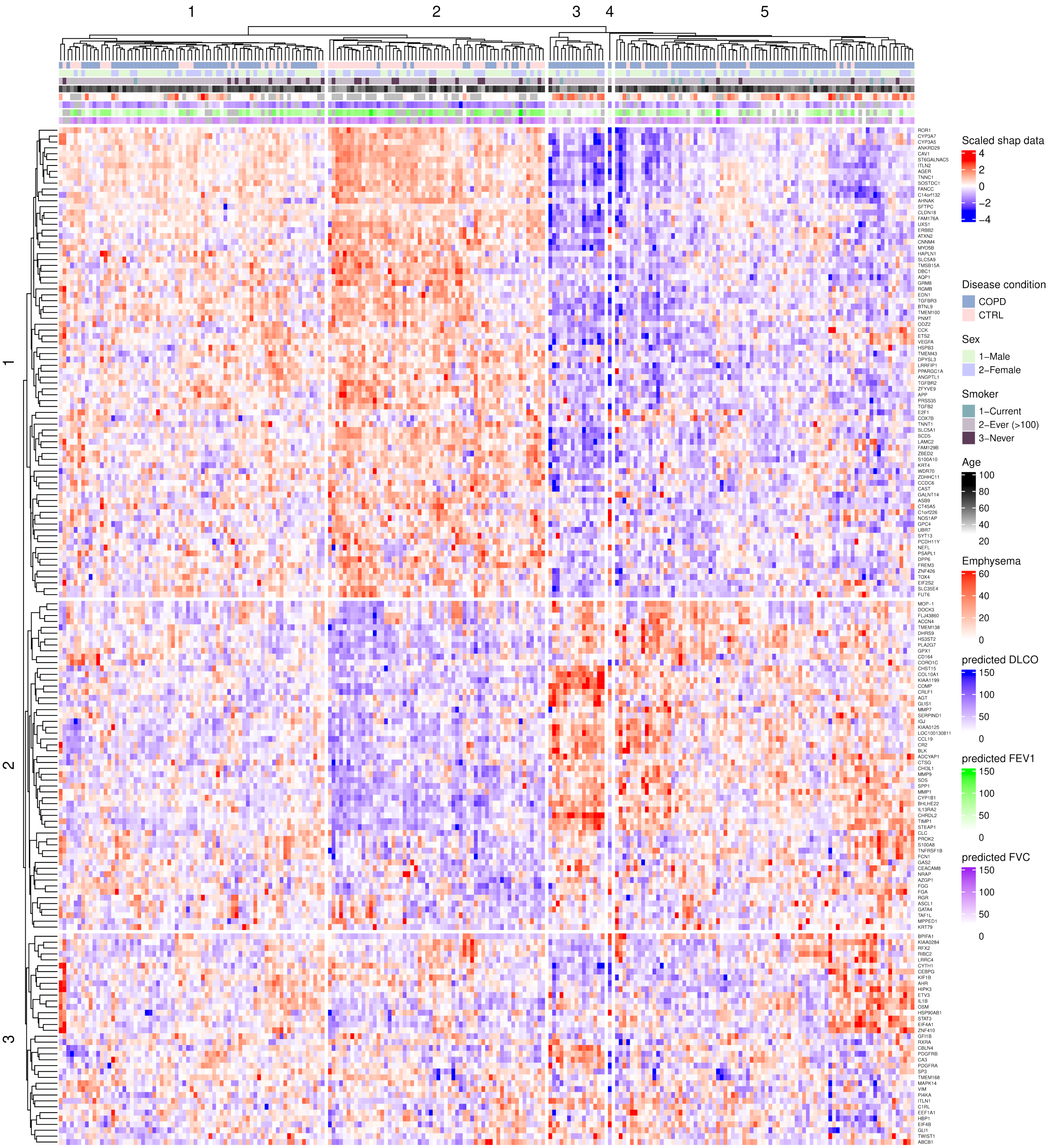


**Fig. S22. Patient clustering based on candidate gene expression.** Heatmap illustrating unsupervised hierarchical clustering of patients (columns) according to the expression profiles of candidate genes (rows). Colour intensity represents scaled gene expression levels. Top annotations indicate clinical variables.

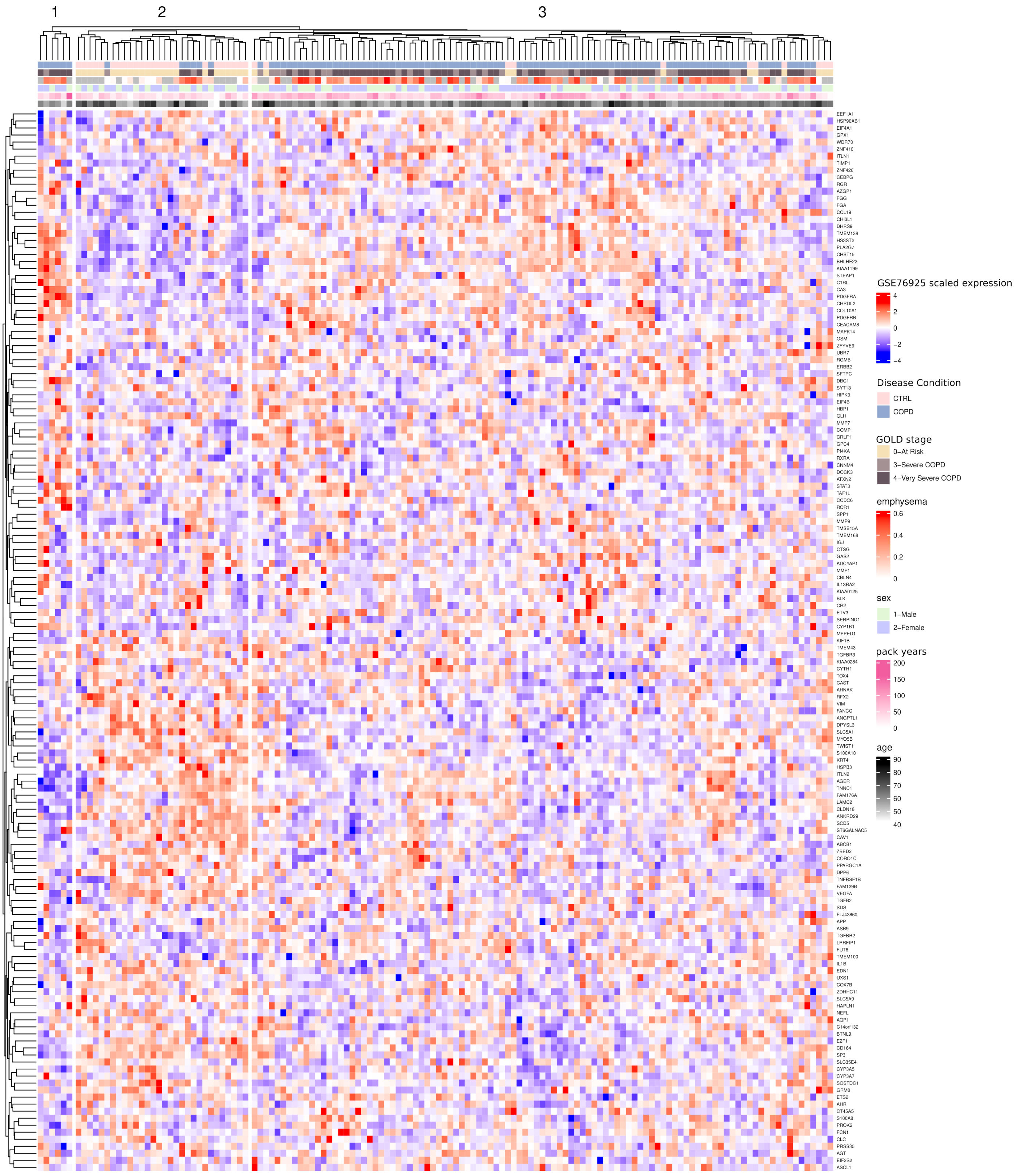


**Fig. S23. Heatmap of scaled expression values for candidate genes in the independent validation cohort GSE76925.** Rows correspond to genes and columns to samples. Hierarchical clustering was performed in an unsupervised manner on both dimensions. Top annotations display clinical variables.

#### 3. SUPPLEMENTARY TABLES

**Table S1.** Collection of tuned hyperparameters of ML models.

| Models | Tuning hyperparameters |
|---|---|
| RF | mtry (number of randomly selected predictors) |
| | min_n (minimal node size) |
| SVM-rad | cost (cost of predicting a sample within or on the wrong side of the margin) |
| | rbf_sigma (radial basis function) |
| SVM-poly | cost (cost of predicting a sample within or on the wrong side of the margin) |
| | degree (polynomial degree) |
| GLM | penalty (amount of regularization) |
| | mixture (proportion of Lasso Penalty) |
| kNN | neighbours (number of neighbours to consider) |
| | dist_power (parameter used in calculating Minkowski distance) |
| | weight_func (type of kernel function used to weight distances between samples) |
| XGB | tree_depth (tree depth) |
| | min_n (minimal node size) |
| | loss_reduction (minimum loss reduction) |
| | sample_size (proportion observations sampled) |
| | mtry (number of randomly selected predictors) |
| | learn_rate (learning rate) |

**Table S2. Summary of classifier performance.** Statistics of normMCC (mean, standard deviation [SD], and median) are provided for both cross-validation (and test) scenarios across all input gene sets. Maximum and minimum values for each scenario are marked in bold.

| Classifier | Mean Value | Standard Deviation (SD) | Median |
|---|---|---|---|
| GLM | 0.738 (0.667) | 0.123 (0.151) | 0.774 (0.720) |
| RF | 0.728 (0.687) | 0.126 (0.139) | 0.762 (0.725) |
| SVM-poly | **0.745** (**0.699**) | 0.129 (**0.168**) | **0.781** (0.720) |
| SVM-rad | 0.739 (0.688) | **0.120** (0.142) | **0.781** (**0.731**) |
| XGB | **0.722** (0.683) | 0.127 (**0.136**) | 0.762 (0.728) |
| kNN | 0.731 (**0.659**) | **0.141** (0.167) | **0.743** **(0.689)** |

Table S3: **Ranked list of candidate genes based on aggregated max normSHAP (Relevance score).** Scores are maintained at 5 decimal places. The list is organised into three parallel blocks to optimise space.

| Rk | Gene | Score | Ct | Rk | Gene | Score | Ct | Rk | Gene | Score | Ct |
|---|---|---|---|---|---|---|---|---|---|---|---|
| 1 | FANCC | 0.18450 | 9 | 59 | GRM8 | 0.02324 | 27 | 116 | CBLN4 | 0.01682 | 30 |
| 2 | KIAA0284 | 0.08886 | 26 | 60 | CD164 | 0.02312 | 27 | 117 | AGT | 0.01679 | 20 |
| 3 | DPYSL3 | 0.07273 | 29 | 61 | TAF1L | 0.02295 | 8 | 118 | SLC5A9 | 0.01666 | 23 |
| 4 | ZBED2 | 0.06489 | 26 | 62 | TNNC1 | 0.02281 | 4 | 119 | SCD5 | 0.01647 | 25 |
| 5 | ROR1 | 0.06439 | 29 | 63 | MAPK14 | 0.02273 | 19 | 120 | AHNAK | 0.01640 | 26 |
| 6 | WDR70 | 0.06353 | 29 | 64 | VIM | 0.02263 | 26 | 121 | LOC100130811 | 0.01634 | 22 |
| 7 | LAMC2 | 0.06200 | 26 | 65 | CR2 | 0.02255 | 28 | 122 | COL10A1 | 0.01619 | 23 |
| 8 | ITLN1 | 0.06114 | 27 | 66 | BHLHE22 | 0.02246 | 30 | 123 | CTSG | 0.01618 | 22 |
| 9 | CYTH1 | 0.06010 | 29 | 67 | AZGP1 | 0.02183 | 26 | 124 | SLC35E4 | 0.01611 | 26 |
| 10 | COX7B | 0.05791 | 28 | 68 | BPIFA1 | 0.02176 | 25 | 125 | ERBB2 | 0.01596 | 16 |
| 11 | GALNT14 | 0.05791 | 29 | 69 | AQP1 | 0.02169 | 35 | 126 | SERPIND1 | 0.01574 | 25 |
| 12 | KRT4 | 0.05389 | 27 | 70 | MMP7 | 0.02156 | 20 | 127 | CHI3L1 | 0.01563 | 28 |
| 13 | ZDHHC11 | 0.05196 | 27 | 71 | DOCK3 | 0.02156 | 7 | 128 | BLK | 0.01540 | 23 |
| 14 | FCN1 | 0.05085 | 34 | 72 | S100A10 | 0.02141 | 27 | 129 | KIF1B | 0.01529 | 25 |
| 15 | CA3 | 0.05001 | 32 | 73 | SDS | 0.02123 | 25 | 130 | TWIST1 | 0.01519 | 21 |
| 16 | CAV1 | 0.04794 | 21 | 74 | MOP-1 | 0.02122 | 24 | 131 | CCDC6 | 0.01514 | 28 |
| 17 | PPARGC1A | 0.04458 | 39 | 75 | CLDN18 | 0.02121 | 24 | 132 | RIBC2 | 0.01498 | 9 |
| 18 | ST6GALNAC5 | 0.04417 | 27 | 76 | PRSS35 | 0.02112 | 26 | 133 | TGFBR2 | 0.01488 | 16 |
| 19 | CRLF1 | 0.04399 | 27 | 77 | IL1B | 0.02110 | 29 | 134 | APP | 0.01483 | 21 |
| 20 | AGER | 0.04270 | 26 | 78 | NOS1AP | 0.02092 | 28 | 135 | GLIS1 | 0.01483 | 25 |
| 21 | C1RL | 0.04240 | 6 | 79 | ZFYVE9 | 0.02073 | 24 | 136 | ADCYAP1 | 0.01470 | 20 |
| 22 | KIAA1199 | 0.04135 | 34 | 80 | PDGFRB | 0.02066 | 19 | 137 | BTNL9 | 0.01469 | 21 |
| 23 | ASCL1 | 0.03962 | 9 | 81 | SOSTDC1 | 0.02064 | 21 | 138 | NRAP | 0.01390 | 28 |
| 24 | TMEM168 | 0.03852 | 5 | 82 | CHRDL2 | 0.02045 | 23 | 139 | E2F1 | 0.01389 | 17 |
| 25 | CYP3A7 | 0.03786 | 23 | 83 | HIPK3 | 0.02043 | 17 | 140 | FAM129B | 0.01381 | 25 |
| 26 | C1orf226 | 0.03760 | 28 | 84 | COMP | 0.02040 | 25 | 141 | GATA4 | 0.01364 | 19 |
| 27 | SPP1 | 0.03581 | 35 | 85 | CCK | 0.02004 | 21 | 142 | IL13RA2 | 0.01360 | 21 |
| 28 | HSPB3 | 0.03527 | 32 | 86 | TMEM100 | 0.01997 | 14 | 143 | EIF2S2 | 0.01359 | 30 |
| 29 | EDN1 | 0.03387 | 27 | 87 | FGG | 0.01995 | 31 | 144 | ATXN2 | 0.01348 | 22 |
| 30 | LRRFIP1 | 0.03175 | 9 | 88 | TNFRSF1B | 0.01994 | 18 | 145 | LRRC4 | 0.01318 | 6 |
| 31 | ZNF426 | 0.03164 | 27 | 89 | FLJ43860 | 0.01993 | 17 | 146 | TIMP1 | 0.01313 | 17 |
| 32 | PI4KA | 0.03144 | 6 | 90 | PROK2 | 0.01973 | 20 | 147 | ODZ2 | 0.01311 | 25 |
| 33 | STAT3 | 0.03067 | 18 | 91 | DPP6 | 0.01966 | 27 | 148 | UBR7 | 0.01298 | 27 |
| 34 | TMEM138 | 0.03010 | 28 | 92 | CNNM4 | 0.01944 | 28 | 149 | RGMB | 0.01297 | 19 |
| 35 | IGJ | 0.03010 | 24 | 93 | MMP1 | 0.01943 | 33 | 150 | GPX1 | 0.01293 | 11 |
| 36 | CT45A5 | 0.02985 | 24 | 94 | HBP1 | 0.01927 | 16 | 151 | OSM | 0.01287 | 22 |
| 37 | CCL19 | 0.02960 | 21 | 95 | TMEM43 | 0.01922 | 25 | 152 | RXRA | 0.01287 | 16 |
| 38 | ASB9 | 0.02719 | 29 | 96 | UXS1 | 0.01920 | 27 | 153 | VEGFA | 0.01286 | 12 |
| 39 | SYT13 | 0.02707 | 32 | 97 | EIF4B | 0.01906 | 24 | 154 | ETV3 | 0.01278 | 18 |
| 40 | CYP1B1 | 0.02678 | 35 | 98 | PLA2G7 | 0.01899 | 28 | 155 | TNNT1 | 0.01248 | 26 |
| 41 | RGR | 0.02677 | 27 | 99 | HAPLN1 | 0.01867 | 32 | 156 | FUT6 | 0.01247 | 24 |
| 42 | SFTPC | 0.02674 | 3 | 100 | S100A8 | 0.01866 | 19 | 157 | FGA | 0.01214 | 9 |
| 43 | TMSB15A | 0.02670 | 25 | 101 | GPC4 | 0.01864 | 29 | 158 | MPPED1 | 0.01206 | 5 |
| 44 | KRT79 | 0.02662 | 27 | 102 | ITLN2 | 0.01862 | 21 | 159 | PCDH11Y | 0.01200 | 25 |
| 45 | ABCB1 | 0.02649 | 20 | 103 | PNMT | 0.01794 | 23 | 160 | CEACAM8 | 0.01197 | 21 |
| 46 | ANKRD29 | 0.02608 | 23 | 104 | ZNF410 | 0.01788 | 16 | 161 | GFI1B | 0.01195 | 18 |
| 47 | GAS2 | 0.02604 | 28 | 105 | HS3ST2 | 0.01787 | 23 | 162 | EIF4A1 | 0.01192 | 5 |
| 48 | SLC5A1 | 0.02600 | 25 | 106 | PDGFRA | 0.01782 | 18 | 163 | CORO1C | 0.01190 | 5 |
| 49 | ACCN4 | 0.02599 | 28 | 107 | FAM176A | 0.01776 | 27 | 164 | TGFB2 | 0.01170 | 28 |
| 50 | KIAA0125 | 0.02591 | 7 | 108 | EEF1A1 | 0.01764 | 20 | 165 | C14orf132 | 0.01149 | 19 |
| 51 | TOX4 | 0.02590 | 27 | 109 | GLI1 | 0.01758 | 17 | 166 | ETS2 | 0.01146 | 18 |
| 52 | CAST | 0.02589 | 25 | 110 | CLC | 0.01750 | 23 | 167 | NEFL | 0.01145 | 22 |

Continued on next page

Table S3: **Ranked list of candidate genes based on aggregated max normSHAP (Relevance score).** Scores are maintained at 5 decimal places. The list is organised into three parallel blocks to optimise space. (Continued)

| | | | | | | | | | | | |
|---|---|---|---|---|---|---|---|---|---|---|---|
| 53 | DHRS9 | 0.02534 | 25 | 111 | FREM3 | 0.01714 | 30 | 168 | CEBPG | 0.01142 | 17 |
| 54 | HSP90AB1 | 0.02483 | 15 | 112 | RFX2 | 0.01697 | 22 | 169 | PSAPL1 | 0.01137 | 25 |
| 55 | CHST15 | 0.02434 | 27 | 113 | DBC1 | 0.01686 | 20 | 170 | MMP9 | 0.01113 | 19 |
| 56 | ANGPTL1 | 0.02393 | 29 | 114 | SP3 | 0.01685 | 20 | 171 | AHR | 0.01105 | 19 |
| 57 | TGFBR3 | 0.02355 | 19 | 115 | MYO5B | 0.01683 | 24 | 172 | CYP3A5 | 0.01097 | 3 |
| 58 | STEAP1 | 0.02325 | 29 | | | | | | | | |

*Note*: Rk: Rank; Score: Relevance score; Ct: Occurrence Count.

**Table S4. Clinical enrichment analysis of patient clusters derived from candidate gene expression.** Statistical evaluation of the association between expression-based patient clusters and clinical variables. For each variable, the statistical test performed, test statistic, and corresponding p-value are reported. Enriched Class indicates the clinical categories significantly overrepresented within each cluster, and cluster denotes the corresponding patient group.

| Variable | Test | Statistic | p-value | Enriched Class | Cluster |
|---|---|---|---|---|---|
| Disease Condition | Chi-squared | 27.878 | $1.292 \times 10^{-7}$ | COPD | 5 |
| GOLD stage | Chi-squared | 34.737 | $4.998 \times 10^{-4}$ | 1-Mild, 2-Mod, 3-Severe, 4-Very Sev | |
| smoker | Chi-squared | 8.576 | $1.499 \times 10^{-2}$ | 1-Current, 2-Ever (>100) | |
| Pred. FEV1 | Wilcoxon | 4968.000 | $7.256 \times 10^{-5}$ | – | |
| emphysema | Wilcoxon | 2411.000 | $1.625 \times 10^{-4}$ | – | |
| Pred. DLCO | Wilcoxon | 6788.500 | $1.168 \times 10^{-7}$ | – | |
| Pred. FVC | T-test | 2.674 | $8.639 \times 10^{-3}$ | – | |
| Disease Condition | Chi-squared | 6.360 | $1.167 \times 10^{-2}$ | COPD | 3 |
| GOLD stage | Chi-squared | 19.291 | $3.998 \times 10^{-3}$ | 3-Severe, 4-Very Severe | |
| Pred. FEV1 | Wilcoxon | 1761.000 | $1.516 \times 10^{-4}$ | – | |
| emphysema | Wilcoxon | 369.000 | $1.129 \times 10^{-4}$ | – | |
| Pred. DLCO | Wilcoxon | 2068.500 | $3.814 \times 10^{-5}$ | – | |
| Pred. FVC | Wilcoxon | 1597.000 | $4.013 \times 10^{-3}$ | – | |
| Disease Condition | Chi-squared | 73.486 | $1.013 \times 10^{-17}$ | CTRL | 2 |
| GOLD stage | Chi-squared | 80.633 | $4.998 \times 10^{-4}$ | 0-At Risk | |
| smoker | Chi-squared | 9.978 | $7.496 \times 10^{-3}$ | 3-Never | |
| Pred. FEV1 | Wilcoxon | 721.500 | $1.040 \times 10^{-13}$ | – | |
| emphysema | Wilcoxon | 3349.500 | $2.980 \times 10^{-7}$ | – | |
| Pred. DLCO | Wilcoxon | 1362.500 | $7.946 \times 10^{-13}$ | – | |
| Pred. FVC | T-test | -5.655 | $2.297 \times 10^{-7}$ | – | |

Table S5: **Association between candidate gene expression and clinical variables.** Adjusted p-values are reported to 3 decimal places. Values $\leq 0.050$ are highlighted in bold.

| Gene | Sex | Smoker | Age | Emphysema | DLCO | FEV1 | FVC |
|---|---|---|---|---|---|---|---|
| FANCC | 1.000 | 0.960 | 0.999 | 0.822 | 0.602 | 0.817 | 0.952 |
| KIAA0284 | 0.826 | 0.955 | 0.999 | 0.929 | 0.769 | 0.894 | 0.952 |
| DPYSL3 | 0.898 | 0.960 | 0.999 | 0.960 | 0.942 | 0.765 | 0.952 |

Continued on next page

Table S5: **Association between candidate gene expression and clinical variables.** Adjusted p-values are reported to 3 decimal places. Values $\leq$ 0.050 are highlighted in bold. (Continued)

| Gene | Sex | Smoker | Age | Emphysema | DLCO | FEV1 | FVC |
|---|---|---|---|---|---|---|---|
| ZBED2 | 0.833 | 0.960 | 0.960 | 0.887 | 0.957 | 0.960 | 0.952 |
| ROR1 | 0.833 | 0.899 | 0.999 | 0.827 | 0.769 | 0.888 | 0.952 |
| WDR70 | 1.000 | 0.854 | 0.999 | 0.644 | 0.917 | 0.960 | 0.952 |
| LAMC2 | 0.826 | 0.854 | 0.727 | 0.887 | 0.917 | 0.960 | 0.952 |
| ITLN1 | 1.000 | 0.854 | 0.777 | 0.887 | 0.767 | 0.960 | 0.975 |
| CYTH1 | 0.826 | 0.960 | 0.536 | 0.927 | 0.893 | 0.968 | 0.952 |
| COX7B | 0.826 | 0.960 | 0.999 | 0.827 | 0.786 | 0.960 | 0.952 |
| GALNT14 | 1.000 | 0.905 | 0.999 | 0.974 | 0.838 | 0.960 | 0.952 |
| KRT4 | 1.000 | 0.960 | 0.999 | 0.874 | 0.319 | 0.655 | 0.952 |
| ZDHHC11 | 0.833 | 0.960 | 0.999 | 0.618 | 0.277 | 0.894 | 0.952 |
| FCN1 | 0.833 | 0.960 | 0.999 | 0.983 | 0.982 | 0.960 | 0.953 |
| CA3 | 1.000 | 0.960 | 0.777 | 0.618 | 0.151 | 0.960 | 0.952 |
| CAV1 | 1.000 | 0.899 | 0.999 | 0.960 | 0.944 | 0.960 | 0.952 |
| PPARGC1A | 0.826 | 0.960 | 0.870 | 0.916 | 0.917 | 0.895 | 0.952 |
| ST6GALNAC5 | 0.826 | 0.960 | 0.171 | 0.979 | 0.774 | 0.811 | 0.952 |
| CRLF1 | 1.000 | 0.960 | 0.999 | 0.980 | 0.957 | 0.966 | 0.952 |
| AGER | 0.826 | 0.955 | 0.999 | 0.827 | 0.597 | 0.960 | 0.952 |
| C1RL | 1.000 | 0.960 | 0.777 | 0.688 | 0.957 | 0.672 | 0.952 |
| KIAA1199 | 0.826 | 0.960 | 0.999 | 0.188 | 0.104 | 0.894 | 0.952 |
| ASCL1 | 1.000 | 0.960 | 0.777 | 0.440 | 0.917 | 0.811 | 0.952 |
| TMEM168 | 0.833 | 0.960 | 0.999 | 0.998 | 0.769 | 0.960 | 0.952 |
| CYP3A7 | 1.000 | 0.854 | 0.999 | 0.887 | 0.769 | 0.811 | 0.952 |
| C1orf226 | 0.826 | 0.960 | 0.999 | 0.916 | 0.917 | 0.983 | 0.953 |
| SPP1 | 1.000 | 0.960 | 0.999 | 0.781 | 0.796 | 0.960 | 0.952 |
| HSPB3 | 0.826 | 0.826 | 0.999 | 0.205 | 0.638 | 0.765 | 0.952 |
| EDN1 | 0.826 | 0.960 | 0.999 | 0.998 | 0.919 | 0.655 | 0.952 |
| LRRFIP1 | 0.833 | 0.854 | 0.999 | 0.644 | 0.420 | 0.960 | 0.952 |
| ZNF426 | 0.826 | 0.854 | 0.921 | 0.998 | 0.957 | 0.960 | 0.952 |
| PI4KA | 0.993 | 0.960 | 0.999 | 0.070 | 0.086 | 0.173 | 0.952 |
| STAT3 | 1.000 | 0.960 | 0.777 | 0.424 | 0.619 | 0.960 | 0.953 |
| TMEM138 | 1.000 | 0.854 | 0.999 | 0.188 | **0.006** | 0.106 | 0.952 |
| IGJ | 0.984 | 0.854 | 0.999 | 0.874 | 0.769 | 0.960 | 0.952 |
| CT45A5 | 1.000 | 0.960 | 0.888 | 0.854 | 0.332 | 0.822 | 0.953 |
| CCL19 | 0.826 | 0.955 | 0.999 | 0.963 | 0.767 | 0.960 | 0.953 |
| ASB9 | 0.826 | 0.960 | 0.794 | 0.644 | 0.984 | 0.820 | 0.952 |
| SYT13 | 1.000 | 0.960 | 0.999 | 0.874 | 0.917 | 0.960 | 0.953 |
| CYP1B1 | 0.833 | 0.854 | 0.796 | 0.434 | 0.966 | 0.960 | 0.952 |
| RGR | 1.000 | 0.854 | 0.999 | 0.729 | 0.838 | 0.960 | 0.952 |
| SFTPC | 0.972 | 0.960 | 0.273 | 0.054 | **0.000** | 0.106 | 0.319 |
| TMSB15A | 0.826 | 0.955 | 0.888 | 0.723 | 0.086 | 0.199 | 0.880 |
| KRT79 | 0.826 | 1.000 | 0.999 | 0.644 | 0.735 | 0.960 | 0.975 |
| ABCB1 | 1.000 | 0.960 | 0.999 | 0.822 | 0.786 | 0.672 | 0.952 |
| ANKRD29 | 1.000 | 0.905 | 0.777 | 0.992 | 0.966 | 0.895 | 0.952 |
| GAS2 | 1.000 | 0.984 | 0.999 | 0.644 | 0.277 | **0.047** | 0.148 |
| SLC5A1 | 1.000 | 0.960 | 0.777 | 0.916 | 0.769 | 0.960 | 0.952 |
| ACCN4 | 0.826 | 0.854 | 0.999 | 0.178 | 0.290 | 0.960 | 0.953 |
| KIAA0125 | 0.993 | 0.854 | 0.999 | 0.455 | 0.917 | 0.655 | 0.952 |
| TOX4 | 0.833 | 0.960 | 0.921 | 0.960 | 0.917 | 0.983 | 0.952 |
| CAST | 0.826 | 0.955 | 0.999 | **0.048** | **0.000** | **0.005** | 0.148 |
| DHRS9 | 0.972 | 0.899 | 0.999 | 0.874 | 0.420 | 0.960 | 0.952 |
| HSP90AB1 | 0.826 | 0.960 | 0.999 | 0.932 | 0.957 | 0.928 | 0.952 |
| CHST15 | 1.000 | 0.960 | 0.777 | 0.188 | 0.088 | 0.630 | 0.952 |

Continued on next page

Table S5: **Association between candidate gene expression and clinical variables.** Adjusted p-values are reported to 3 decimal places. Values ≤ 0.050 are highlighted in bold. (Continued)

| Gene | Sex | Smoker | Age | Emphysema | DLCO | FEV1 | FVC |
|---|---|---|---|---|---|---|---|
| ANGPTL1 | 0.826 | 0.854 | 0.999 | 0.644 | 0.367 | 0.526 | 0.120 |
| TGFBR3 | 1.000 | 0.960 | 0.999 | 0.827 | 0.942 | 0.995 | 0.952 |
| STEAP1 | 0.826 | 0.960 | 0.999 | 0.822 | 0.332 | 0.928 | 0.952 |
| GRM8 | 1.000 | 0.899 | 0.999 | 0.963 | 0.917 | 0.983 | 0.952 |
| CD164 | 1.000 | 0.960 | 0.999 | 0.178 | 0.102 | **0.031** | 0.066 |
| TAF1L | 1.000 | 0.854 | 0.999 | 0.874 | 0.957 | 0.888 | 0.953 |
| TNNC1 | 0.826 | 0.960 | 0.999 | 0.827 | 0.917 | 0.960 | 0.952 |
| MAPK14 | 1.000 | 0.899 | 0.921 | 0.715 | 0.919 | 0.737 | 0.952 |
| VIM | 1.000 | 0.899 | 0.999 | 0.960 | 0.917 | 0.811 | 0.553 |
| CR2 | 0.898 | 0.960 | 0.999 | 0.054 | 0.290 | 0.863 | 0.953 |
| BHLHE22 | 1.000 | 0.960 | 0.999 | 0.470 | 0.213 | 0.655 | 0.952 |
| AZGP1 | 1.000 | 0.960 | 0.601 | 0.960 | 0.786 | 0.655 | 0.952 |
| BPIFA1 | 1.000 | 0.960 | 0.999 | 0.451 | 0.096 | 0.672 | 0.952 |
| AQP1 | 1.000 | 0.999 | 0.999 | 0.820 | 0.838 | 0.960 | 0.952 |
| MMP7 | 0.826 | 0.960 | 0.999 | 0.916 | 0.919 | 0.655 | 0.952 |
| DOCK3 | 1.000 | 0.960 | 0.999 | 0.424 | 0.225 | 0.655 | 0.952 |
| S100A10 | 1.000 | 0.955 | 0.920 | 0.054 | **0.001** | 0.121 | 0.494 |
| SDS | 1.000 | 0.960 | 0.999 | 0.874 | 0.917 | 0.960 | 0.952 |
| MOP-1 | 0.826 | 0.960 | 0.999 | 0.744 | 0.350 | 0.863 | 0.952 |
| CLDN18 | 1.000 | 0.960 | 0.395 | 0.477 | 0.235 | 0.227 | 0.553 |
| PRSS35 | 0.759 | 0.960 | 0.395 | 0.612 | 0.138 | 0.200 | 0.952 |
| IL1B | 1.000 | 0.960 | 0.999 | 0.822 | 0.865 | 0.960 | 0.952 |
| NOS1AP | 0.833 | 0.960 | 0.999 | 0.451 | 0.638 | 0.995 | 0.952 |
| ZFYVE9 | 1.000 | 0.955 | 0.999 | 0.960 | 0.823 | 0.960 | 0.952 |
| PDGFRB | 0.833 | 0.960 | 0.999 | 0.822 | 0.435 | 0.960 | 0.952 |
| SOSTDC1 | 1.000 | 0.960 | 0.777 | 0.512 | 0.917 | 0.960 | 0.953 |
| CHRDL2 | 1.000 | 0.960 | 0.999 | 0.612 | 0.769 | 0.894 | 0.952 |
| HIPK3 | 0.826 | 0.960 | 0.921 | 0.916 | 0.786 | 0.928 | 0.953 |
| COMP | 0.826 | 0.960 | 0.999 | 0.688 | 0.735 | 0.895 | 0.953 |
| CCK | 1.000 | 0.854 | 0.888 | 0.941 | 0.823 | 0.960 | 0.952 |
| TMEM100 | 0.972 | 0.986 | 0.999 | 0.644 | 0.235 | 0.655 | 0.953 |
| FGG | 1.000 | 0.999 | 0.999 | 0.960 | 0.907 | 0.894 | 0.952 |
| TNFRSF1B | 1.000 | 0.960 | 0.777 | 0.992 | 0.966 | 0.948 | 0.952 |
| FLJ43860 | 1.000 | 0.854 | 0.999 | 0.434 | 0.332 | 0.960 | 0.953 |
| PROK2 | 0.826 | 0.854 | 0.999 | 0.644 | 0.765 | 0.960 | 0.952 |
| DPP6 | 1.000 | 0.960 | 0.999 | 0.932 | 0.942 | 0.928 | 0.952 |
| CNNM4 | 1.000 | 0.960 | 0.999 | 0.983 | 0.957 | 0.928 | 0.952 |
| MMP1 | 0.833 | 0.960 | 0.951 | 0.927 | 0.676 | 0.655 | 0.952 |
| HBP1 | 1.000 | 0.960 | 0.999 | 0.887 | 0.332 | 0.672 | 0.952 |
| TMEM43 | 1.000 | 0.955 | 0.999 | 0.932 | 0.448 | 0.960 | 0.952 |
| UXS1 | 1.000 | 0.999 | 0.999 | 0.190 | 0.786 | 0.960 | 0.952 |
| EIF4B | 1.000 | 0.854 | 0.999 | 0.887 | 0.917 | 0.960 | 0.952 |
| PLA2G7 | 0.972 | 0.960 | 0.999 | 0.960 | 0.178 | 0.695 | 0.952 |
| HAPLN1 | 0.910 | 0.955 | 0.999 | 0.434 | 0.950 | 0.630 | 0.952 |
| S100A8 | 1.000 | 0.960 | 0.999 | 0.916 | 0.769 | 0.960 | 0.978 |
| GPC4 | 1.000 | 0.960 | 0.999 | 0.827 | 0.769 | 0.524 | 0.553 |
| ITLN2 | 1.000 | 0.960 | 0.999 | 0.956 | 0.919 | 0.894 | 0.952 |
| PNMT | 1.000 | 0.960 | 0.888 | 0.891 | 0.917 | 0.985 | 0.952 |
| ZNF410 | 1.000 | 0.960 | 0.999 | 0.874 | 0.917 | 0.983 | 0.952 |
| HS3ST2 | 1.000 | 0.960 | 0.999 | 0.887 | 0.602 | 0.960 | 0.952 |
| PDGFRA | 1.000 | 0.854 | 0.251 | **0.048** | 0.315 | 0.173 | 0.952 |
| FAM176A | 0.826 | 0.923 | 0.395 | 0.792 | 0.786 | 0.960 | 0.952 |

Continued on next page

Table S5: **Association between candidate gene expression and clinical variables.** Adjusted p-values are reported to 3 decimal places. Values $\leq$ 0.050 are highlighted in bold. (Continued)

| Gene | Sex | Smoker | Age | Emphysema | DLCO | FEV1 | FVC |
|---|---|---|---|---|---|---|---|
| EEF1A1 | 1.000 | 0.984 | 0.888 | 0.916 | 0.435 | 0.812 | 0.952 |
| GLI1 | 0.826 | 0.960 | 0.708 | 0.283 | 0.444 | 0.672 | 0.952 |
| CLC | 1.000 | 0.960 | 0.999 | 0.814 | 0.618 | 0.121 | 0.148 |
| FREM3 | 1.000 | 0.960 | 0.960 | 0.960 | 0.917 | 0.960 | 0.952 |
| RFX2 | 0.993 | 0.960 | 0.999 | 0.983 | 0.736 | 0.960 | 0.953 |
| DBC1 | 1.000 | 0.960 | 0.999 | 0.054 | 0.306 | 0.655 | 0.952 |
| SP3 | 1.000 | 0.854 | 0.999 | 0.887 | 0.917 | 0.978 | 0.953 |
| MYO5B | 0.826 | 0.970 | 0.999 | 0.644 | 0.315 | 0.725 | 0.952 |
| CBLN4 | 1.000 | 0.960 | 0.870 | 0.556 | 0.917 | 0.655 | 0.952 |
| AGT | 1.000 | 0.960 | 0.324 | 0.188 | 0.907 | 0.121 | 0.066 |
| SLC5A9 | 0.972 | 0.905 | 0.999 | 0.979 | 0.368 | 0.983 | 0.952 |
| SCD5 | 1.000 | 0.955 | 0.999 | 0.603 | 0.420 | 0.121 | 0.467 |
| AHNAK | 0.972 | 0.960 | 0.164 | 0.874 | 0.957 | 0.960 | 0.952 |
| LOC100130811 | 0.826 | 0.960 | 0.999 | 0.916 | 0.767 | 0.960 | 0.953 |
| COL10A1 | 1.000 | 0.854 | 0.171 | 0.170 | 0.225 | 0.655 | 0.978 |
| CTSG | 0.833 | 0.960 | 0.251 | 0.527 | 0.602 | 0.655 | 0.953 |
| SLC35E4 | 0.904 | 0.960 | 0.999 | 0.832 | 0.917 | 0.957 | 0.953 |
| ERBB2 | 1.000 | 0.955 | 0.923 | 0.960 | 0.960 | 0.960 | 0.978 |
| SERPIND1 | 0.833 | 0.960 | 0.999 | 0.916 | 0.319 | 0.812 | 0.952 |
| CHI3L1 | 0.972 | 0.955 | 0.999 | 0.960 | 0.957 | 0.983 | 0.978 |
| BLK | 1.000 | 0.984 | 0.999 | 0.960 | 0.367 | 0.811 | 0.952 |
| KIF1B | 0.826 | 0.960 | 0.921 | 0.998 | 0.850 | 0.928 | 0.952 |
| TWIST1 | 0.826 | 0.960 | 0.963 | 0.999 | 0.957 | 0.895 | 0.953 |
| CCDC6 | 0.826 | 0.960 | 0.907 | 0.792 | 0.561 | 0.983 | 0.953 |
| RIBC2 | 0.898 | 0.960 | 0.395 | 0.998 | 0.917 | 0.960 | 0.952 |
| TGFBR2 | 1.000 | 0.960 | 0.999 | 0.960 | 0.982 | 0.960 | 0.952 |
| APP | 0.864 | 0.998 | 0.888 | 0.190 | 0.638 | 0.817 | 0.953 |
| GLIS1 | 0.833 | 0.960 | 0.357 | 0.105 | **0.009** | 0.226 | 0.952 |
| ADCYAP1 | 0.826 | 0.960 | 0.999 | 0.874 | 0.786 | 0.672 | 0.953 |
| BTNL9 | 1.000 | 0.955 | 0.999 | 0.983 | 0.957 | 0.960 | 0.952 |
| NRAP | 1.000 | 0.960 | 0.777 | 0.188 | 0.151 | 0.655 | 0.952 |
| E2F1 | 0.826 | 0.955 | 0.921 | 0.884 | 0.786 | 0.822 | 0.952 |
| FAM129B | 1.000 | 0.960 | 0.779 | 0.190 | 0.917 | 0.121 | 0.953 |
| GATA4 | 1.000 | 0.960 | 0.796 | 0.147 | 0.638 | 0.655 | 0.952 |
| IL13RA2 | 1.000 | 0.854 | 0.777 | 0.874 | 0.638 | 0.655 | 0.952 |
| EIF2S2 | 1.000 | 0.960 | 0.999 | 0.922 | 0.769 | 0.655 | 0.952 |
| ATXN2 | 0.826 | 0.998 | 0.999 | 0.960 | 0.774 | 0.960 | 0.952 |
| LRRC4 | 0.833 | 0.960 | 0.999 | 0.927 | 0.420 | 0.928 | 0.952 |
| TIMP1 | 1.000 | 0.960 | 0.888 | 0.575 | 0.638 | 0.655 | 0.952 |
| ODZ2 | 0.898 | 0.955 | 0.999 | 0.916 | 0.917 | 0.795 | 0.952 |
| UBR7 | 0.833 | 0.854 | 0.322 | 0.644 | 0.206 | 0.755 | 0.953 |
| RGMB | 1.000 | 0.960 | 0.822 | 0.887 | 0.917 | 0.983 | 0.953 |
| GPX1 | 1.000 | 0.905 | 0.999 | 0.832 | 0.895 | 0.928 | 0.953 |
| OSM | 1.000 | 0.960 | 0.418 | 0.729 | 0.823 | 0.960 | 0.953 |
| RXRA | 1.000 | 0.960 | 0.999 | 0.910 | 0.769 | 0.894 | 0.953 |
| VEGFA | 0.833 | 0.854 | 0.171 | 0.054 | **0.006** | 0.173 | 0.952 |
| ETV3 | 1.000 | 0.960 | 0.888 | 0.744 | 0.638 | 0.655 | 0.952 |
| TNNT1 | 1.000 | 0.960 | 0.999 | 0.716 | 0.769 | 0.765 | 0.952 |
| FUT6 | 1.000 | 0.999 | 0.999 | 0.386 | 0.700 | 0.655 | 0.953 |
| FGA | 1.000 | 0.960 | 0.999 | 0.960 | 0.213 | 0.817 | 0.952 |
| MPPED1 | 1.000 | 0.999 | 0.999 | 0.952 | 0.865 | 0.672 | 0.953 |
| PCDH11Y | 0.826 | 0.999 | 0.999 | 0.916 | 0.602 | 0.894 | 0.952 |

Continued on next page

Table S5: **Association between candidate gene expression and clinical variables.** Adjusted p-values are reported to 3 decimal places. Values $\leq 0.050$ are highlighted in bold. (Continued)

| Gene | Sex | Smoker | Age | Emphysema | DLCO | FEV1 | FVC |
|---|---|---|---|---|---|---|---|
| CEACAM8 | 1.000 | 0.960 | 0.999 | 0.982 | 0.917 | 0.894 | 0.952 |
| GFI1B | 0.972 | 0.955 | 0.999 | 0.917 | 0.917 | 0.655 | 0.952 |
| EIF4A1 | 0.865 | 0.999 | 0.999 | 0.963 | 0.769 | 0.655 | 0.952 |
| CORO1C | 0.972 | 0.960 | 0.305 | 0.095 | **0.002** | 0.121 | 0.198 |
| TGFB2 | 0.993 | 0.960 | 0.999 | 0.960 | 0.917 | 0.666 | 0.952 |
| C14orf132 | 0.993 | 0.960 | 0.923 | 0.714 | 0.769 | 0.960 | 0.952 |
| ETS2 | 1.000 | 0.998 | 0.999 | 0.992 | 0.982 | 0.948 | 0.952 |
| NEFL | 0.833 | 0.960 | 0.777 | 0.512 | 0.301 | 0.173 | 0.952 |
| CEBPG | 0.972 | 0.960 | 0.777 | 0.512 | 0.786 | 0.655 | 0.952 |
| PSAPL1 | 1.000 | 0.960 | 0.999 | 0.188 | 0.602 | 0.894 | 0.952 |
| MMP9 | 0.833 | 0.960 | 0.999 | 0.822 | 0.826 | 0.960 | 0.952 |
| AHR | 0.898 | 0.854 | 0.999 | 0.644 | 0.805 | 0.960 | 0.953 |
| CYP3A5 | 0.993 | 0.960 | 0.305 | 0.188 | 0.235 | 0.672 | 0.952 |

*Note*: DLCO: Predicted DLCO; FEV1: Predicted FEV1; FVC: Predicted FVC.

**Table S6. External validation performance of ML classifiers using candidate genes.** Reported values correspond to predictive performance (normMCC) for RF, kNN, SVM-rad, SVM-pol, GLM, and XGB across independent datasets. For GSE47460, the main value represents cross-validation performance, while the value in parentheses corresponds to the test set performance after retraining only on the candidate gene list.

| Dataset | RF | kNN | SVM-rad | SVM-pol | GLM | XGB |
|---|---|---|---|---|---|---|
| GSE47460 (this study) | 0.82 (0.75) | 0.86 (0.83) | 0.85 (0.75) | 0.85 (0.76) | 0.84 (0.74) | 0.83 (0.79) |
| GSE8581 – GSE37768 | 0.67 | 0.62 | 0.65 | 0.65 | 0.65 | 0.58 |
| GSE69818 – GSE103174 | 0.60 | 0.64 | 0.63 | 0.64 | 0.64 | 0.57 |
| GSE76925 | 0.53 | 0.61 | 0.46 | 0.45 | 0.45 | 0.56 |

**Table S7. Overlap between candidate genes and DEA genes across external datasets.** Results of enrichment analyses comparing candidate genes with DEA genes in independent lung tissue datasets. For each dataset, the FDR and $\log_2$FC thresholds used for DEA are indicated. The table also shows the observed overlap with candidate genes and the expected overlap from Monte Carlo permutation analysis.

| Dataset | FDR | $\log_2$FC | DEA genes | Observed overlap (cand. $\cap$ DEA) | Expected overlap (permutations) |
|---|---|---|---|---|---|
| GSE57148 (RNA-seq) | 0.05 | 0.5 | 821 | 13 | 0.04 |
| GSE76925 | 0.05 | 0 | 3091 | 44 | 0 |
| | 0.05 | 0.25 | 712 | 24 | 0 |
| | 0.05 | 0.5 | 53 | 11 | 0 |
| | 0.01 | 0 | 1710 | 25 | $9 \times 10^{-4}$ |
| | 0.01 | 0.25 | 627 | 16 | 0 |
| | 0.01 | 0.5 | 46 | 7 | 0 |

**Table S8. Comparison of ML performance with previously published microarray tissue-based models.** Cross-validation and test set metrics for sensitivity, specificity, and normMCC are reported. This table highlights the high predictive performance of our best models and the robustness of the selected candidate gene set relative to existing approaches.

| Regulation Study | Cross-validation Set | | | Test Set | | |
|---|---|---|---|---|---|---|
| | **Sensitivity** | **Specificity** | **normMCC** | **Sensitivity** | **Specificity** | **normMCC** |
| This study – best model | 0.97 (svm-p mRMR) | 0.86 (glm data-driven) | 0.87 (knn mRMR) | 1.00 | 0.80 | 0.81 |
| This study – final candidate | 0.87 (knn) | 0.88 (svm-rad) | 0.87 (knn) | 0.92 | 0.72 | 0.83 |
| Gohari – Lasso (2023) | 0.93 | 0.81 | – | 0.83 | 0.44 | – |
| Gohari – SCAD (2023) | 0.87 | 0.80 | – | 0.75 | 0.67 | – |
| Gohari – MCP (2023) | 0.90 | 0.79 | – | – | – | – |
| Mostafaei (2018) | 0.85 | 0.51 | – | – | – | – |
| Yao (2019) | 0.87 | 0.81 | – | 0.93 | 0.87 | – |
